# De novo Design of a Peptide Modulator to Reverse Sodium Channel Dysfunction Linked to Cardiac Arrhythmias and Epilepsy

**DOI:** 10.1101/2025.04.25.649609

**Authors:** Ryan Mahling, Bence Hegyi, Erin Cullen, Timothy M. Cho, Aaron Rodriques, Lucile Fossier, Marc Yehya, Lin Yang, Bi-Xing Chen, Alexander N. Katchman, Nourdine Chakouri, Ruiping Ji, Elaine Wan, Jared Kushner, Steven O. Marx, Sergey Ovchinnikov, Christopher Makinson, Donald M. Bers, Manu Ben-Johny

**Affiliations:** Department of Physiology and Cellular Biophysics, Columbia University, New York, NY, USA; Department of Pharmacology, University of California Davis, Davis, CA, USA; Department of Neurology, Columbia University, New York, NY, USA; Division of Cardiology, Department of Medicine, Columbia University, New York, NY, USA; Department of Pharmacology, Columbia University, New York, NY, USA; John Harvard Distinguished Science Fellowship Program, Harvard University, Cambridge, MA, USA; Department of Neuroscience, Columbia University, New York, NY, USA

## Abstract

Ion channels orchestrate electrical signaling in excitable cells. In nature, ion channel function is customized by modulatory proteins that have evolved to fulfill distinct physiological needs. Yet, engineering synthetic modulators that precisely tune ion channel function is challenging. One example involves the voltage-gated sodium (Na_V_) channel that initiates the action potential, and whose dysfunction amplifies late/persistent sodium current (*I*_NaL_), a commonality that underlies various human diseases including cardiac arrhythmias and epilepsy. Here, using a computational protein design platform, we engineered a *de novo* peptide modulator, ELIXIR, that binds Na_V_ channels with submicromolar affinity. Functional analysis revealed an unexpected selectivity in inhibiting ‘pathogenic’ *I*_NaL_ and confirmed its effectiveness in reversing Na_V_ dysfunction linked to both cardiac arrhythmias and epilepsy in cellular and murine models. These findings exemplify the efficacy of *de novo* protein design for engineering synthetic ion channel modulators and sets the stage for rational design of future therapeutic approaches.

## Introduction

Ion channels are pore-forming proteins that serve as nature’s transistor^1^. Unlike their silicon counterparts, ion channel activity is dynamic and tightly regulated, allowing cells to adapt their electrical properties to match the complex requirements of life. Indeed, ion channel dysfunction underlies life-threatening human diseases including cardiac arrhythmias^2-4^ and epilepsy^5^, and well as myotonia^6^, migraines^7^, and intellectual disability^8^. For many voltage-gated ion channels, pathophysiological changes in channel function result in altered channel dynamics that disrupt overall electrical signaling. As such, strategies to precisely tune ion channel function are highly desired; yet current approaches, including small-molecule pharmacology, are limited and often have off-target effects^9,10^. In nature, the functional plasticity of ion channels, a defining feature of these proteins, is often attained through association with diverse auxiliary subunits or regulatory proteins that have evolved to fine tune specific channel properties^11^. Structural studies over the past several decades have furnished an unprecedented atomistic view of such interactions and shed light onto mechanisms of regulation^12-15^. Yet, rational engineering of *de* novo or synthetic ion channel modulators with high specificity has remained intractable.

A prominent example of this deficiency involves the family of voltage-gated sodium channels (Na_V_1.1-1.9) which are responsible for the generation and propagation of action potentials (APs) in excitable cells, including cardiomyocytes and neurons. A hallmark of Na_V_ channel function is the fast activation and inactivation that drives the rapid depolarization of many excitable cells, including neurons and muscle cells. Disruption of Na_V_ channel function is linked to a broad spectrum of clinical phenotypes, including cardiac arrhythmias, congenital myotonia, epilepsy, neurodevelopmental disorders and painful neuropathies. Despite their distinct etiology, many of these disorders are linked to deficits in channel inactivation that can contribute to sustained Na^+^ influx, referred to as either late or persistent Na^+^ current (*I*_NaL_). Two prominent examples involve the *SCN5A* gene, which encodes Na_V_1.5 in the heart, and the *SCN8A* gene, which encodes Na_V_1.6 in the nervous system. Specifically, gain-of-function (GoF) human variants in Na_V_1.5 that upregulate *I*_NaL_ are linked to arrhythmogenic long-QT syndrome 3 (LQT3)^16^, and atrial fibrillation^2^. Similarly, pathological remodeling associated with heart failure (HF)^3^ or following myocardial ischemia^17^, has been shown to modify Na_V_1.5 function and elevate *I*_NaL_, resulting in an increased risk of cardiac arrhythmias. Increased *I*_NaL_ causes a proarrhythmic prolongation of the AP^4,18,19^ (**Figure 1A**) and disrupts Na^+^ and Ca^2+^ homeostasis, which sets off a vicious and detrimental cycle involving CaMKII, ROS, and RyR^20^. Similarly, in the nervous system, GoF Na_V_1.6 variants are linked to severe developmental epilepsies and sudden unexpected death in epilepsy (SUDEP)^21^. These variants disrupt sodium channel inactivation and alter neuronal action potential properties, to ultimately drive neuronal network hyperexcitability.

**Figure 1.**
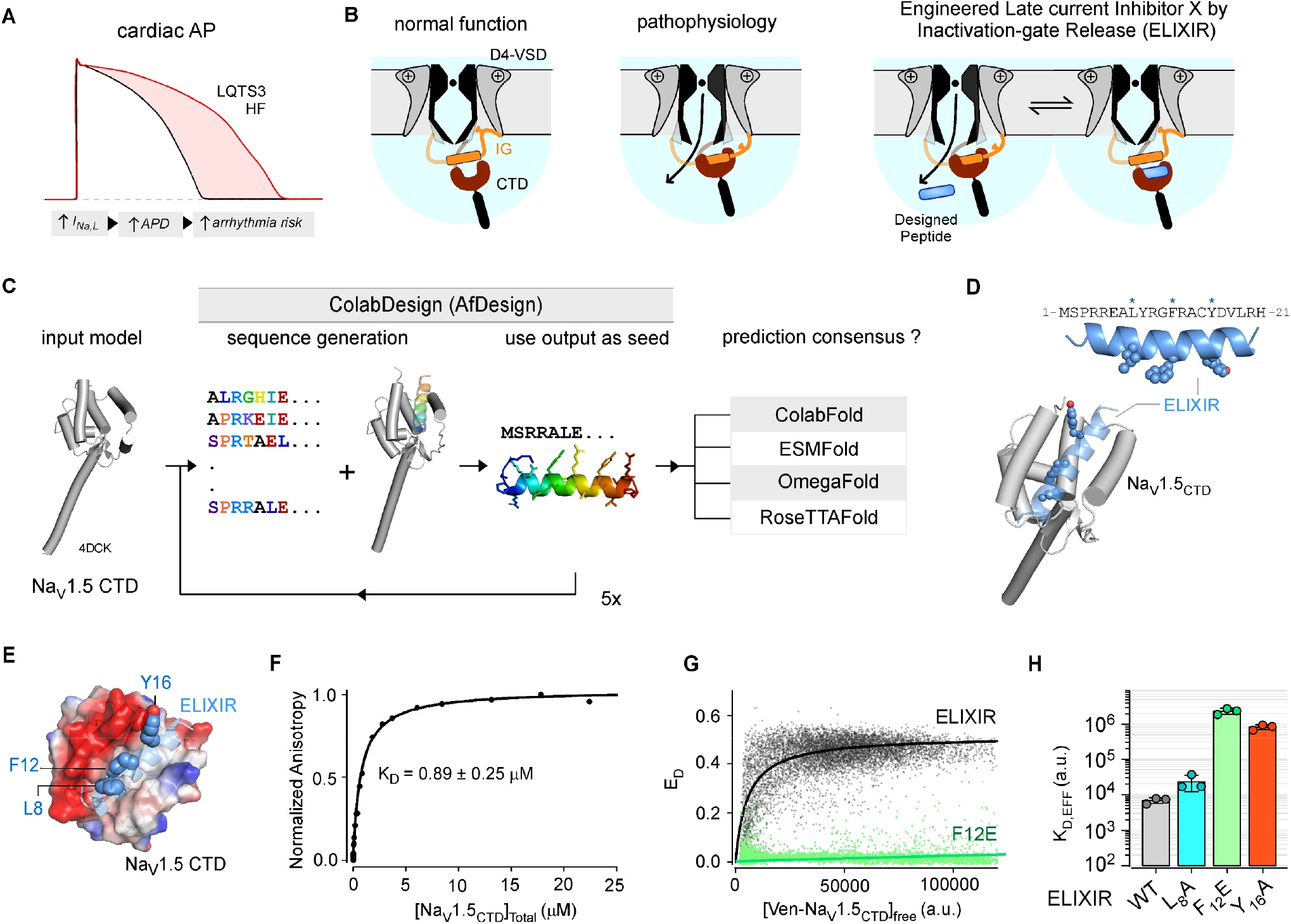
*De novo* design of a peptide modulator of Na_V_1.5. **(A)** Increased I_NaL_ prolongs AP duration and increases the risk of cardiac arrhythmias. **(B)** Conceptual scheme for rational design of an *I*_NaL_ inhibitor. Left, following channel activation, the inactivation gate (IG) binds to a pore-proximal site to inactivate Na_V_1.5 (black/gray; IG, orange; EFL, dark red). Middle, in pathophysiology, IG has reduced efficacy for binding the pore-proximal site and remains associated to CTD. Competitive displacement of IG from EFL by the designed peptide named ELIXIR may promote inactivation and inhibit *I*_NaL_. **(C)** Schematic shows the peptide design framework. The initial coordinates of the Na_V_1.5 CTD (PDBID: 4DCK) served as input into AfDesign to hallucinate a peptide binder (rainbow helix). Structural predictions of ELIXIR–Na_V_1.5 CTD complex were compared to assess consensus. **(D)** Top, helical model of ELIXIR along with protein sequence. Residues L8, F12, and Y16 are shown as ball and stick in the helical model. Bottom, ColabFold predicted model of the Na_V_1.5 CTD - ELIXIR complex. **(E)** Detailed view of the ColabFold predicted Na_V_1.5 CTD EFL bound to ELIXIR. ELIXIR residues L8, F12, and Y16 are shown as ball and stick. Vacuum electrostatic surface (positive; blue, neutral, white; negative, red). **(F)** Equilibrium titration of ELIXIR with the Na_V_1.5_CTD_ show saturable binding. Average *K*_D_ and standard deviation were obtained from 3 replicate titrations. **(G)** FRET efficiency between Venus-tagged Na_V_1.5_CTD_ and Cerulean-tagged ELIXIR (black) or ELIXIR F12E mutant plotted against the concentration of free Venus-tagged Na_V_1.5_CTD_, a.u. is arbitrary units. Compared to wild-type ELIXIR, F12E mutant displays weak FRET. The solid line depicts the fit of a 1:1 binding isotherm. **(H)** Bar graph summary of *K*_d,EFF_, the relative dissociation constant for ELIXIR-Na_V_1.5_CTD_ interaction obtained from FRET 2-hybrid experiments. ELIXIR mutations variably disrupt binding.

Beyond cardiac and epileptic disorders, increased *I*_NaL_ through related channels expressed in skeletal muscle or the peripheral nervous system is associated with congenital myotonia^6^ and pain disorders^22^, pointing to the potential utility of *I*_NaL_ inhibitors. Despite advances in the understanding of pathophysiological mechanisms and the availability of high-resolution structures^23,24^, devising effective strategies to reverse pathophysiological changes in Na_V_ function has been limited. The prevailing approach has been small molecule *I*_NaL_ blockers such as Ranolazine, which show promise. However, these drugs primarily target a common antiarrhythmic drug binding site in the transmembrane region, which may variably inhibit peak Na^+^ current or block other ion channels^9^. In addition, these small molecules may also have reduced affinity for disease-linked mutant channels, limiting their potential utility. Recently, certain auxiliary proteins, such as the intracellular fibroblast growth factor homologous factors (iFGF/FHF), have been shown to inhibit *I*_NaL_ with minimal effects on peak Na current^25,26^. Additionally, peptides derived from iFGF/FHF have been shown to be sufficient to recapitulate these effects^26^. Even still, despite optimization, these peptides are comparatively large and show reduced efficacy for certain channel variants. To this end, major advances in machine-learning based computational protein design have increased the accuracy of *de novo* protein design while simultaneously reducing computational time^27-29^. These emerging methods may provide an alternative strategy to rationally design peptide modulators that precisely modify Na_V_ function. If substantiated, these approaches may be generalized to engineer a toolkit of synthetic ion channel regulators.

With cardiac Na_V_1.5 as a template, we used *de novo* protein design to generate a peptide modulator that selectively inhibits pathogenic *I*_NaL_ by enhancing a native mechanism of channel inactivation. To do so, we implemented a protocol for peptide hallucination within the ColabDesign framework^30,31^ and leveraged this to generate a short 21-aa peptide targeting the Na_V_1.5 carboxy-terminal domain (CTD), an intracellular channel domain that serves as a regulatory hub. *In vitro* binding analysis confirmed the interaction of the candidate peptide with the Na_V_1.5 CTD. In depth electrophysiological analysis confirmed the functionality of the *de novo* designed peptide and revealed an unexpected selectivity of the designed peptide for pathogenic *I*_NaL_ of multiple members of the Na_V_ channel family. We further demonstrate the effectiveness of this peptide in both neuronal and cardiac disease models. Our findings highlight the utility of *de novo* protein design in the development of synthetic ion channel modulators that tune specific aspects of ion channel function in physiology and pathophysiology.

## Results

### A rational approach to developing an *I*_NaL_ inhibitor with submicromolar affinity

Fast inactivation of Na_V_ channels is a multistep process involving distinct channel elements. Following depolarization, movement of voltage-sensing domains (VSDs) I-III couples to pore opening, while that of the domain IV VSD initiates conformational changes that result in inactivation^32-34^. Of particular importance for the latter is the inactivation gate (IG, linker between domains III and IV), which contains the IFM motif that binds to a site near the channel pore to halt Na^+^ influx^35,36^ (**Figure 1B**). In addition to this pore-proximal site, the IG also binds an acidic EF-hand like region (EFL) in the C-terminal domain (CTD) of the channel^37-40^, suggesting that the IG may first need to be released from this EFL site to interact with the pore-proximal site and prevent pore opening (**Figure S1A**). In cardiac pathologies, this process is disrupted resulting in decreased Na_V_1.5 inactivation and increased *I*_NaL_^4,19^, suggesting reduced occupancy of the pore-proximal site by the IG (**Figure S1B**). Given this, we reasoned that dislodging the IG from its CTD interaction site would promote Na_V_ inactivation, and in so doing, reduce *I*_NaL_ and AP duration (APD, **Figure 1B, S1C**). To do so, we used a ColabDesign^31^ based protocol to generate a peptide targeting the EFL of the Na_V_1.5 CTD as depicted in **Figure 1C**. This process yielded a small (21 aa) peptide with features reminiscent of the Na_V_1.5 IG (hydrophobic surface flanked by basic residues, consistent with it being able to bind the hydrophobic cleft in the EFL of the Na_V_1.5 CTD (**Figure 1D-E**). Given its putative mechanism, we termed this peptide, *Engineered Late-current* <u>*I*</u>*nhibitor* <u>*X*</u> *by* <u>*I*</u>*nactivation-gate Release* (ELIXIR). As an initial means of testing our designed sequence we compared models of the ELIXIR-Na_V_1.5 CTD (aa 1773 – 1940, Na_V_1.5_CTD_) complex generated with several structure prediction algorithms^27,41-43^ (**Figure S2A**). In all models, the Na_V_1.5 EFL and IQ motif adopted a nearly identical architecture (backbone RMSD ∼ 1.58A) to that observed in crystallographic structures^44-46^ (**Figure S2A**), lending credence to the predicted structures. In each, ELIXIR binds to the Na_V_1.5 EFL in an identical orientation. The interface between ELIXIR and Na_V_1.5 EFL was highly similar across the models, with peptide residues L8, F12, and Y16 making numerous close contacts to residues in the hydrophobic cleft of the EFL (**Figure S2A – S2I**). In addition to this, ColabFold models of the Na_V_1.5 CTD-CaM-Na_V_1.5 IG complex in the absence and presence of ELIXIR show ELIXIR bound to the EFL in place of the IG (**Figure S2J**), suggesting it can displace the IG from this site. Comparison of our designed peptide sequence to that of the Na_V_1.5 IG revealed that they are only ∼24% identical (**Figure S3A**), indicating that our designed sequence was not merely recapitulating the native ligand of the Na_V_1.5 EFL. Indeed, when the sequence was searched against the BLAST database^47^, the closest homolog was the 50S ribosome of the algae *Nannochloropsis gaditana* which showed only ∼52% identity (**Figure S3A**). The closest mammalian homolog, having ∼33% identity, was DNA replication factor CDT1 of the Weddell seal (*Leptonychotes weddellii*, **Figure S3A**). A single conservative substitution was made to the sequence (C15S) to simplify *in vitro* applications of ELIXIR. Having confirmed that the ELIXIR sequence was both *de novo* and foreign to Na_V_1.5, we examined whether it binds directly to the channel CTD. Equilibrium titrations of a fluorescein-tagged ELIXIR with the Na_V_1.5_CTD_, monitored through fluorescence anisotropy, displayed saturable binding and revealed that ELIXIR binds the Na_V_1.5_CTD_ *in vitro* with a submicromolar affinity (*K*_D_ = 0.89 ± 0.25 μM, **Figure 1F**). To ensure that ELIXIR can associate with the Na_V_1.5_CTD_ in the complex milieu of living cells, we undertook a flow-cytometry based FRET 2-hybrid assay. Co-expression of Cerulean-tagged ELIXIR, a FRET donor, and Venus-tagged Na_V_1.5_CTD_, a FRET acceptor, resulted in robust FRET confirming that the Na_V_1.5_CTD_ binds ELIXIR in mammalian cells (**Figure 1G**). We sought to further validate the interface between ELIXIR and Na_V_1.5 CTD. As L8, F12, and Y16 of ELIXIR were predicted to make many close contacts with residues in the hydrophobic cleft of the Na_V_1.5_CTD_ EFL (**Figure 1E**), we reasoned that the substitution of these aliphatic residues with alanine (L8 and Y16) or glutamate (F12) would disrupt the ELIXIR-Na_V_1.5 CTD interaction. Consistent with this expectation, FRET 2-hybrid analysis showed that each mutant diminished binding, albeit to different degrees **(Figure 1G-1H, S3B-C)**. Collectively, these results demonstrate that *de novo* designed ELIXIR peptide interacts with Na_V_1.5 CTD and may be positioned to tune channel function.

### ELIXIR decreases *I*_NaL_ of a Na_V_1.5 channelopathic variants

Thus affirmed, we sought to determine the functional consequences of ELIXIR interaction on Na_V_1.5 channel function. Whole-cell patch-clamp recordings demonstrated that co-expression of ELIXIR did not significantly alter channel activation, inactivation, or recovery from inactivation of wild-type Na_V_1.5 (**Figure S3D – S3G**). To probe whether ELIXIR inhibits pathological *I*_NaL_, we conducted cell-attached multichannel recordings of HEK293 cells heterologously expressing either wild-type Na_V_1.5 or a disease-linked non-sense variant that truncates the channel CTD (Δ1864). As the channel unitary current is larger than instrument noise, multichannel recordings allow us to quantify peak current and simultaneously detect rare late channel openings in the context of an intact cell, as established previously^25,26,48^. We quantified *I*_NaL_ as *R*_persist_, the ratio of average open probability (*P*_O_) following 50 ms of depolarization normalized to the peak *P*_O_. As in previous studies^25,26,48^, wild-type Na_V_1.5 exhibited a low baseline level of *I*_NaL_, which was minimally perturbed by co-expression of ELIXIR (**Figure 2A-C, S4A**). By contrast, the Δ1864 mutant showed greatly (∼40-fold) elevated *I*_NaL_ at baseline, consistent with the pathogenicity of this variant (**Figure 2C-E, S4B**). Strikingly, ELIXIR co-expression evoked a greater than 12-fold reduction in *I*_NaL_ of the mutant channel. To ensure that the C15S substitution we introduced into ELIXIR did not affect its efficacy as an *I*_NaL_ inhibitor, we also probed the unmodified sequence (**Figure S4C-D**). Both peptides caused a nearly identical reduction in the *I*_NaL_ of Na_V_1.5 Δ1864 (**Figure S4E**). In native tissue, Na_V_ channels function as multi-subunit complexes that include auxiliary *β*-subunits, which have been reported to modulate *I*_*NaL*_^49,50^. As such, we investigated whether ELIXIR could inhibit *I*_*NaL*_ *in* the presence of *β*-subunits. In the presence of β1 and β4, ELIXIR had minimal effects on the *I*_NaL_ of wild-type Na_V_1.5 (**Figure S4F-J**) However, expression of β1 and β4 with the Δ1864 mutant yielded a baseline reduction in *I*_Na,L_, which was further reduced by the addition of ELIXIR (**Figure S4F-J**). Ranolazine, is a well-established clinically used small molecule *I*_NaL_ inhibitor that interacts with Na_V_1.5 through the antiarrhythmic drug (AAD) binding site within the transmembrane segment^24^. We found that incubation with 10 μM ranolazine caused an ∼4-fold reduction in *I*_NaL_ of Na_V_1.5 Δ1864 (**Figure 2C, 2F, S5A**). These findings establish the biological activity of the computationally designed ELIXIR peptide and highlight its effectiveness.

**Figure 2.**
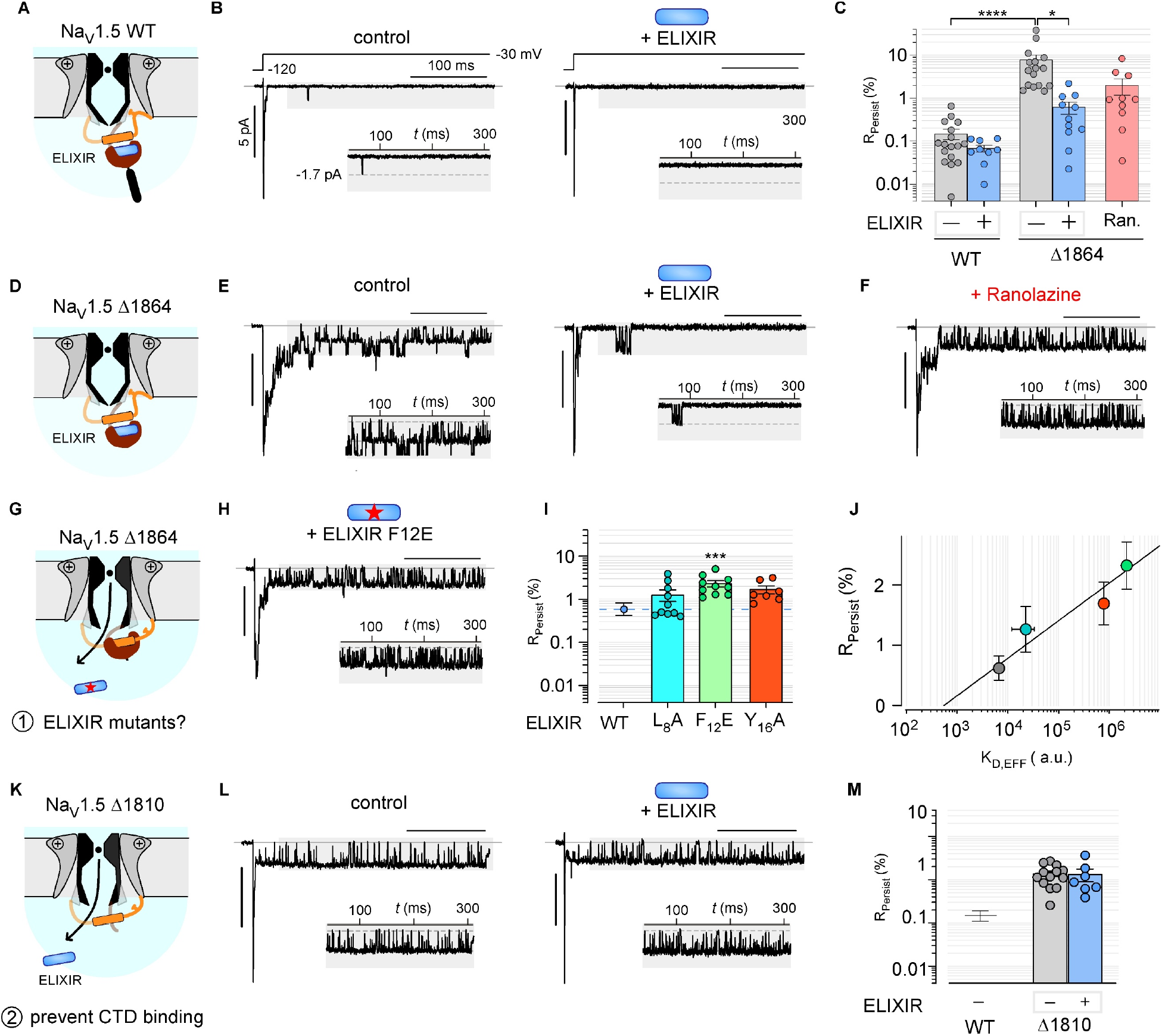
Functional inhibition of *I*_NaL_ by ELIXIR. **(A)** Schematic shows ELIXIR interaction with the CTD of wild-type Na_V_1.5. **(B)** Exemplar multichannel recordings of wild-type Na_V_1.5 in HEK293 cells in the absence (left) and presence of ELIXIR (right). Channel openings are downward deflections from the zero line (gray). Insets show the late phase enlarged for better visualization. **(C)** Bar graph summary shows changes in *I*_*NaL*_ quantified as *R*_persist_. Wild-type Na_V_1.5 without (n = 17; 1069 sweeps) and with ELIXIR (n = 9; 924 sweeps). Na_V_1.5 Δ1864 without ELIXIR (n = 17; 907 sweeps), with ELIXIR (n = 11; 948 sweeps), or with 10 µM ranolazine (n = 10; 848 sweeps). Each bar, mean ± SEM. \*\*\*\**p* < 0.001 compared to wild-type, \**p* = 0.0147 compared to Na_V_1.5 Δ18641 + ELIXIR by a Kruskal-Wallis test followed by Dunn’s multiple comparison test. **(D-F)** ELIXIR inhibits *I*_NaL_ of Na_V_1.5 Δ1864. Schematic (panel D), Exemplar multichannel recordings of Na_V_1.5 Δ1864 mutant in the absence and presence of ELIXIR (panel E), and with Ranolazine (panel F). Format as in Panels A-C. **(G)** Schematic shows weak association of ELIXIR mutant F12E to Na_V_1.5 Δ1864 mutant. **(H)** Exemplar multichannel recordings of Na_V_1.5 Δ1864 mutant with ELIXIR F12E mutant. **(I)** Bar graph summary of changes in R_persist_ of Na_V_1.5 Δ1864 when expressed with ELIXIR L8A (n = 9, 852 sweeps), ELIXIR F12E (n = 10, 885 sweeps), or ELIXIR Y16A (n = 8, 671 sweeps). The grey and light blue dashed lines show the average R_persist_ of Na_V_1.5 Δ1864 alone and co-expressed with ELIXIR, respectively. Each bar, mean ± SEM. \*\*\**p* = 0.0008 (+ELIXIR F12E), compared to Na_V_1.5 Δ1864 in the presence of WT ELIXIR by a Kruskal-Wallis test followed by Dunn’s multiple comparison test. **(J)** The relative affinity of Venus-tagged Na_V_1.5_CTD_ for Cerulean-tagged WT and mutant ELIXIR sequences plotted against the R_persist_ of Na_V_1.5 Δ1864 when co-expressed with the corresponding ELIXIR construct. The trendline between the data points is shown in solid black. **(K)** Schematic shows Na_V_1.5 Δ1810 mutant that lacks the ELIXIR-interacting CTD. **(L)** Exemplar multichannel recordings of Na_V_1.5 Δ1810 mutant with ELIXIR F12E mutant. **(M)** Bar graph summary of changes in R_persist_ of Na_V_1.5 Δ1810 when expressed with ELIXIR.

Having confirmed its functionality, we sought to further investigate whether the change in *I*_NaL_ observed with ELIXIR results from its bona fide interaction with Na_V_1.5 CTD. To do so, we first examined how mutations of key ELIXIR residues that weaken binding to the Na_V_1.5 CTD alter functional inhibition of *I*_NaL_ (**Figure 2G**). We found that F12E strongly diminished *I*_NaL_ inhibition, while Y16A and L8A evoked partial effects (**Figure 2H-I, S5B-S5F**). Quantitative analysis revealed a correlation between the level of *I*_NaL_ (*R*_persist_) and the relative dissociation constant determined by FRET analysis (**Figure 2J**), pointing to the functional importance of the ELIXIR -Na_V_1.5 CTD interaction. Second, we reasoned that deletion of the ELIXIR binding site (i.e. the EFL) would abolish *I*_NaL_ inhibition by ELIXIR. To test this, we considered the channelopathic Na_V_1.5 Δ1810 variant, which encodes only the first helix of the EFL (**Figure 2K**). As with the Δ1864 mutant, Na_V_1.5 Δ1810 showed a markedly increased basal *I*_NaL_ relative to the wild-type channel (**Figure 2L-M, S5G**). However, unlike Na_V_1.5 Δ1864, co-expression of ELIXIR had virtually no effect on *I*_NaL_ (**Figure 2L-M, S5G**). Together, the reduced efficacy of ELIXIR mutants in inhibiting *I*_NaL_, and the inability of ELIXIR to inhibit *I*_NaL_ of Na_V_1.5 Δ1810 suggest that ELIXIR interacts with holo-Na_V_1.5 via an interface similar to that predicted in the models of the ELIXIR-Na_V_1.5_CTD_ complex.

### ELIXIR is a general inhibitor of Na_V_1.5 *I*_NaL_

At the molecular level, a variety of perturbations to Na_V_1.5 contribute to pathogenic increases in *I*_NaL_. These include variants distributed throughout the channel sequence and post-translational modifications, such as phosphorylation, that are associated with cardiac pathologies including heart failure^51^. Our results with Na_V_1.5 Δ1864 suggest that ELIXIR is a potent inhibitor of *I*_NaL_; thus, we sought to determine its generality in tuning *I*_NaL_ triggered by distinct mechanisms. Accordingly, we assessed the efficacy of ELIXIR in inhibiting *I*_NaL_ of several Na_V_1.5 mutations located in distinct channel domains. These included: (1) the ΔKPQ varint linked to LQT3, (2) a mutation (F1759A) in the classic AAD binding pocket linked to atrial fibrillation^2^, and (3) a structure-guided mutation (IQ/AA) engineered to block binding of the auxiliary protein calmodulin (**Figure 3A**). These mutations were selected as they represent disparate structural mechanisms that can contribute to increased *I*_NaL_ in cardiac pathologies. Consistent with our previous results^26^, when expressed alone, each of these Na_V_1.5 mutants displayed an elevated *I*_NaL_ (21-fold F1759A; 13-fold ΔKPQ; 10-fold IQ/AA) relative to the wild-type channel (**Figure 3B – 3C,3E, S6A-S6D**). For each Na_V_1.5 mutant, co-expression of ELIXIR resulted in a statistically significant decrease in *I*_NaL_, with reductions ranging from 12-fold (F1759A) to 3-fold (IQ/AA) (**Figure 3B – 3C,3E, S6A-S6D**).

**Figure 3.**
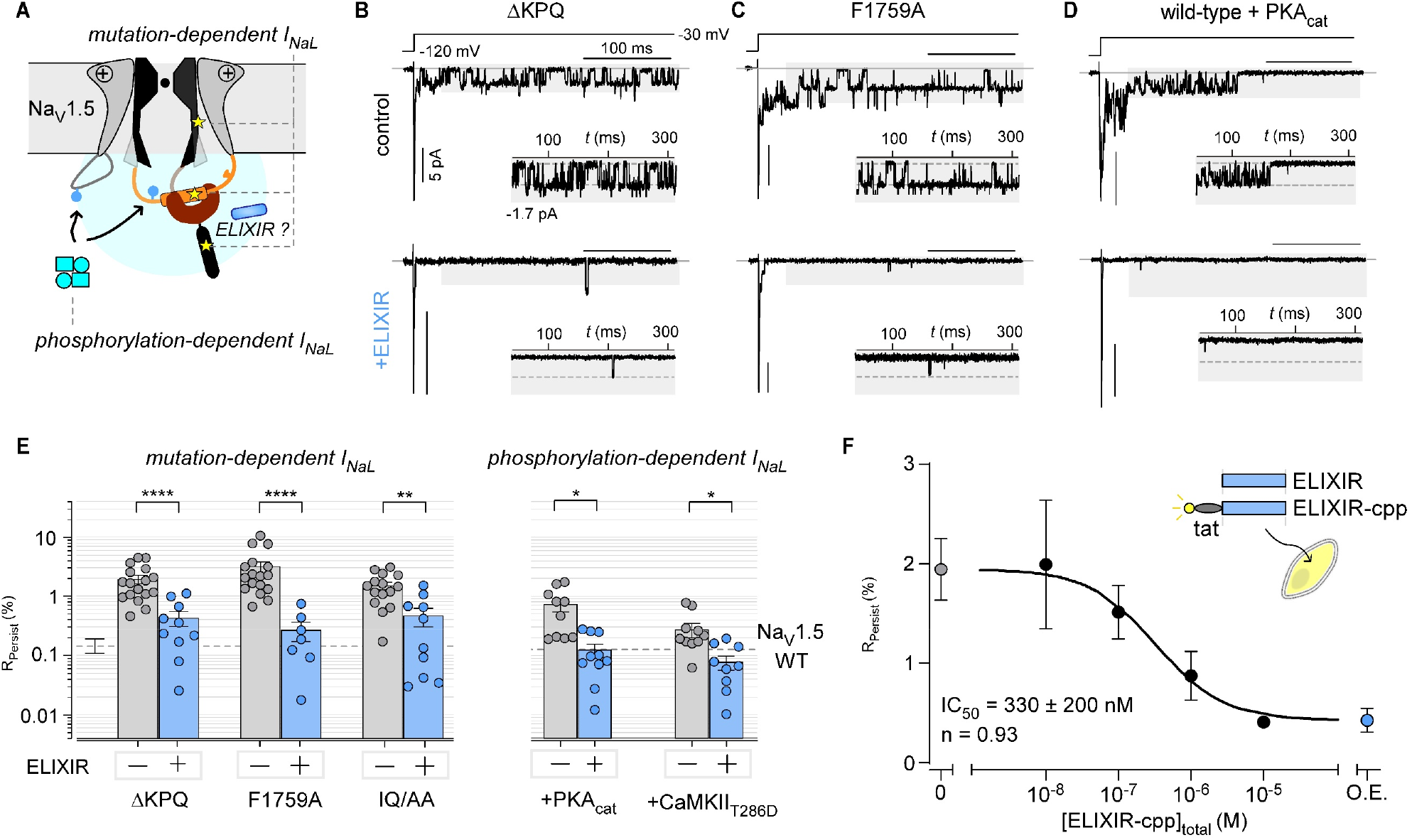
Generality of late Na^+^ current inhibition by ELIXIR. **(A)** Schematic showing that either mutations in or phosphorylation of Na_V_1.5 can lead to increased *I*_NaL_. Stars designate mutations and blue circles phosphorylation. **(B - D)** Exemplar multichannel recordings of Na_V_1.5 ΔKPQ (**B**), Na_V_1.5 F1759A (**C**) and WT Na_V_1.5 with PKA_Cat_ (**D**) in the absence (top) and presence of ELIXIR (bottom). Insets show the late phase enlarged for better visualization. **(E)** Bar graph summarizes effect of ELIXIR on *I*_NaL_ resulting from various mutations and phosphorylation, quantified as *R*_persist_. ΔKPQ without (n = 17, sweeps = 995) and with ELIXIR (n = 10; 936 sweeps); F1759A without (n = 17; 1083 sweeps) and with ELIXIR (n = 7; 725 sweeps); IQ/AA without (n = 15; 926 sweeps) and with ELIXIR (n = 10; 977 sweeps), WT Na_V_1.5 + PKA_Cat_ without (n = 10; 710 sweeps) and with ELIXIR (n = 10; 841 sweeps); Na_V_1.5 + CaMKII_T286D_ without (n = 11; 1033 sweeps) and with +ELIXIR (n = 10; 1029 sweeps). Each bar, mean ± SEM. \*\*\*\**p* < 0.0001 (ΔKPQ, F1759A), \*\**p* = 0.0015 (IQ/AA), \**p* = 0.0101 (+PKA_Cat_), \**p* = 0.0101 (CaMKII_T286D_) by a two-tailed Mann-Whitney U-test. **(F)** Schematic shows ELIXIR modified with a cell-penetrating peptide to facilitate intracellular delivery (ELIXIR cpp). *I*_*NaL*_ of Na_V_1.5 ΔKPQ following treatment with different concentrations of ELIXIR-cpp. The IC_50_ and n values were obtained through a global fit of the data with a Hill equation. The error values of the reported IC_50_ are the 95% confidence interval of the fit. O.E. shows the *I*_*NaL*_ when ELIXIR is overexpressed.

Post-translational modification of Na_V_1.5 also contributes to elevated *I*_NaL_ and is associated with increased risk of arrhythmia in both heart failure^51,52^ and myocardial ischemia^53,54^. To determine whether ELIXIR inhibits phosphorylation-dependent *I*_NaL_, we co-expressed wild-type Na_V_1.5 with the PKA catalytic subunit (PKA_cat_) or a constitutively activated Ca^2+^/CaM-dependent kinase II mutant (CaMKII_T286D_). In both cases, we observed an increase in basal *I*_NaL_ (∼5-fold with PKA_Cat_ or ∼ 2-fold with CaMKII_T286D_) (**Figure 3D-3E, S6E-S6G**). Co-expression of ELIXIR sufficed to reverse the PKA_cat_ and CaMKII_T286D_-dependent enhancement of *I*_NaL_ (**Figure 3D-3E, S6E-S6G**). The ability of ELIXIR to inhibit *I*_NaL_ stemming from a wide range of molecular alterations (i.e., mutations in distinct channel domains and post-translational modifications) indicates that it may be generalizable and further highlights its potential utility.

We next examined the dose-dependence of *I*_NaL_ inhibition by ELIXIR. To facilitate intracellular delivery of ELIXIR, we synthesized it as a peptide fused to the cell-penetrating region of the transduction domain of HIV-1 trans activator of transcription and an N-terminal fluorescein (ELIXIR-cpp). We incubated HEK293 cells transfected with Na_V_1.5 ΔKPQ with various concentrations of ELIXIR-cpp for 4 hours and measured *I*_NaL_ (**Figure 3F, S6H-S6K**). We found that ELIXIR decreased the *I*_NaL_ of Na_V_1.5 ΔKPQ in a dose-dependent manner with an IC50 (∼ 330 nM ± 230 nM (95% CI), **Figure 3F**), similar to the *in vitro* affinity of ELIXIR for the Na_V_1.5_CTD_.

### ELIXIR preferentially tunes “pathological” *I*_NaL_

The nine Na_V_ isoforms (1.1 – 1.9) are well known to have distinct inactivation properties that are critical for their physiological functions^55^. GoF variants in multiple Na_V_ isoforms are linked to diverse clinical phenotypes, including epilepsy^21^, myotonia^6^, neurodevelopmental disorders^56^, and chronic pain^22^ (**Figure 4A**). The high homology of the CTD (**Figure 4B**) suggests that ELIXIR may interact with each Na_V_ isoform. Consistent with this, in ColabFold^41^ generated models ELIXIR was predicted to adopt a highly similar binding pose with the EFL of Na_V_1.1 - 1.9 (**Figure S7A-S7H**). Accordingly, we sought to determine the effect of ELIXIR on *I*_NaL_ of selected wild-type Na_V_ channels expressed in distinct tissues: (1) Na_V_1.1 in neurons, (2) Na_V_1.4 in skeletal muscle, (3) Na_V_1.6 in neurons, and (4) Na_V_1.7 in peripheral nervous system neurons. When expressed in HEK293 cells, these channels exhibited varying amplitudes of *I*_NaL_, with *R*_persist_ values ranging from ∼0.5% (Na_V_1.4) – 1.6% (Na_V_1.1). Co-expression of ELIXIR failed to evoke a statistically significant change in the basal *I*_NaL_ of any of the wild-type Na_V_ isoforms examined: Na_V_1.1 (**Figure 4C,E and S7I**), Na_V_1.7 (**Figure 4D-E and S7J)**, Na_V_1.4 (**Figure 4F, 4H and S7K)**, and Na_V_1.6 **(Figure 4I, 4K and S7M**)

**Figure 4.**
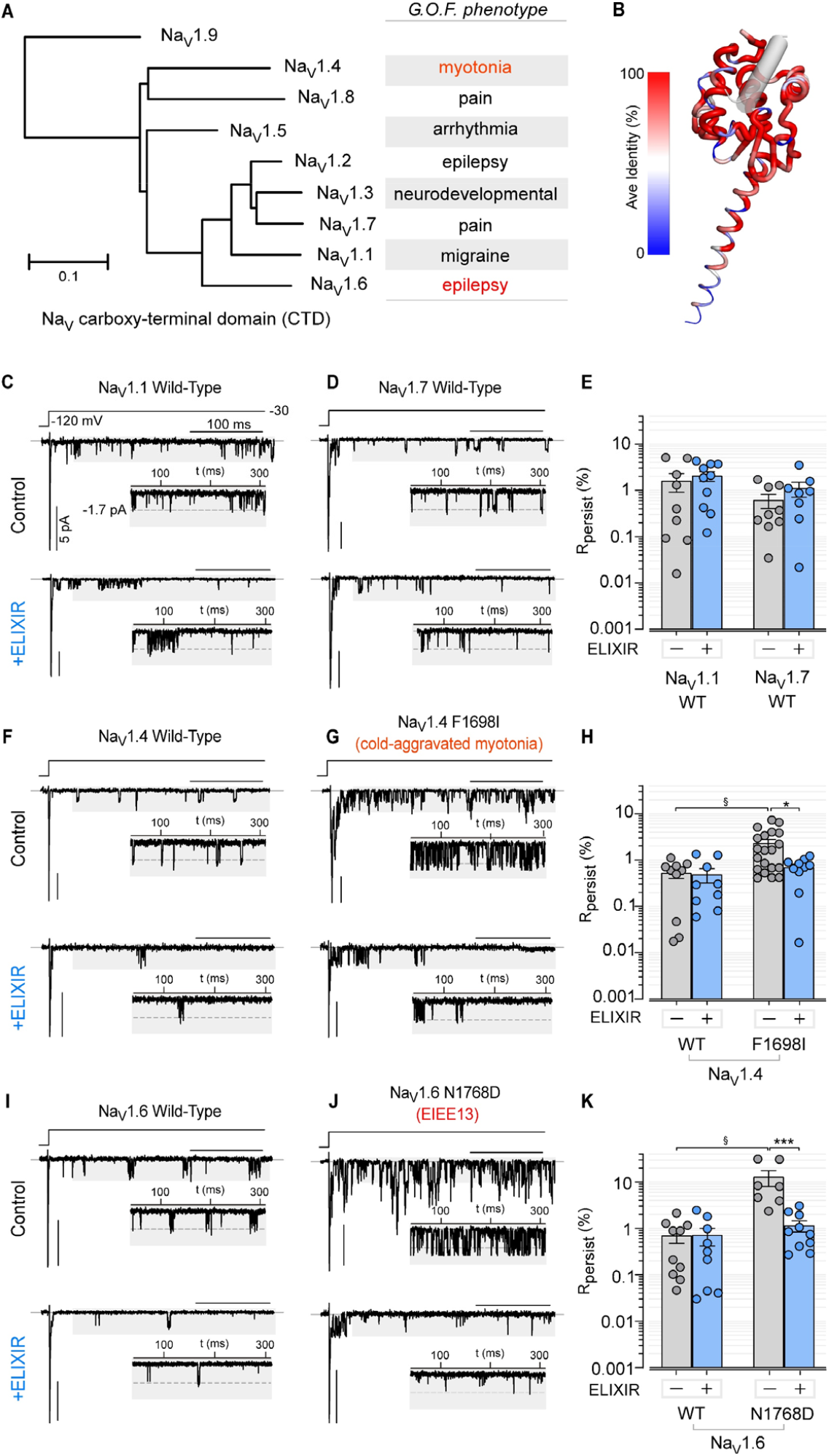
Selectivity of ELIXIR for pathogenic late Na^+^ current. **(A)** Phylogenetic tree of the nine human Na_V_ CTD sequences along with gain of function phenotypes. **(B)** ColabFold model of the Na_V_1.5 CTD+ELIXIR complex. Na_V_1.5 CTD cartoon thickness and color correspond to the average identity to the Na_V_1.5 sequence at each position. ELIXIR is shown as a gray cylinder. **(C – D)** Exemplar multichannel recordings show effect of ELIXIR on wild-type Na_V_1.1 (**C**), Na_V_1.7 (**D**) channels. Top, without ELIXIR; Bottom, with ELIXIR. Insets show the channel openings in the late phase enlarged for better visualization. **(E)** Bar graph summary of effect of ELIXIR on wild-type Na_V_1.1 and Na_V_1.7. Each bar, mean ± SEM, Na_V_1.1 without (n = 9; 912 sweeps) and with ELIXIR (n = 10; 880 sweeps); Na_V_1.7 without (n = 9; 833 sweeps) and with ELIXIR (n = 8; 765 sweeps). **(F – G)** Exemplar multichannel recordings show effect of ELIXIR on wild-type (**F**) and F1698I mutant (**G**) Na_V_1.4channels. Top, without ELIXIR; Bottom, with ELIXIR. Insets show the channel openings in the late phase enlarged for better visualization. **(H)** Bar graph summarizes effect of ELIXIR on wild-type and mutant Na_V_1.4 channels. Each bar, mean ± SEM. Na_V_1.4 without (n = 10, 953 sweeps), and with ELIXIR (n = 9, 812 sweeps); Na_V_1.4 F1698I mutant (n = 20; 1647 sweeps) and with ELIXIR (n = 10; 956 sweeps). §*p* = 0.0101 (wild-type Na_V_1.4 and Na_V_1.4 F1698I), \**p* = 0.0118 (Na_V_1.4 F1698I mutant with and without ELIXIR) by a one-way ANOVA followed by a Tukey’s multiple comparisons test. **(I – J)** Exemplar multichannel recordings show effect of ELIXIR on wild-type (**I**) and N1768D mutant (**J**) Na_V_1.6channels. Top, without ELIXIR; Bottom, with ELIXIR. Insets show the channel openings in the late phase enlarged for better visualization. **(K)** Bar graph summarizes effect of ELIXIR on wild-type and mutant Na_V_1.6 channels. Each bar, mean ± SEM. Na_V_1.6 without (n = 10; 973 sweeps) and with ELIXIR (n = 9; 885 sweeps); Na_V_1.6 N1768D mutant without (n = 7; 876 sweeps) and with ELIXIR (n = 10; 1162 sweeps). §*p* = 0.0009 (wild-type Na_V_1.6 and Na_V_1.6 N1768D),, \**p* = 0.0008 (Na_V_1.6 N1768D mutant with and without ELIXIR) by a one-way ANOVA followed by a Tukey’s multiple comparisons test.

Thus affirmed, we probed whether ELIXIR could modulate *I*_NaL_ of mutant channels that have deficits in inactivation (i.e. if the IG has reduced propensity for occupying the pore-proximal site). In this regard, increased *I*_NaL_ of Na_V_1.4 is considered a relevant pathophysiological mechanism for congenital myotonia^6^. To probe this possibility, we investigated the effect of ELIXIR on the *I*_NaL_ of an Na_V_1.4 variant (F1698I) associated with cold-aggravated myotonia in humans^57,58^. When expressed alone, Na_V_1.4 F1698I had a small (∼4-fold) but statistically significant increase in *R*_persist_ compared to wild-type Na_V_1.4 (**Figure 4F-4H, S7K-S7L**). Co-expression of ELIXIR reversed *I*_NaL_ of the mutant channel to near wild-type levels (**Figure 4F-4H, S7L**), despite the wild-type channel being unperturbed by ELIXIR.

Gene variants in voltage-gated sodium channels are a leading cause of monogenic epilepsies^59^. Variants in *SCN8A*, encoding Na_V_1.6, often increase neuronal excitability by impairing inactivation and increasing *I*_NaL_ ^21,60,61^. Consistent with previous findings^21,62-64^, analysis of an Na_V_1.6 variant (N1768D) linked to epilepsy and seizure disorders showed a markedly elevated *I*_NaL_ as compared to the wild-type channel (**Figure 4I-K, S7M-S7N**). Strikingly, we found that co-expression of ELIXIR reduced *I*_NaL_ (∼10-fold) of Na_V_1.6 N1768D to near wild-type levels (**Figure 4I-4K, S7M-S7N**). These findings suggest that although ELIXIR may bind multiple Na_V_ isoforms, its inhibition of *I*_NaL_ is most prominent only when there are defects in the inactivation process. This feature may allow ELIXIR to preferentially modulate mutant or pathologically modified channels which could be advantageous.

### ELIXIR reverses altered neuronal excitability linked to Na_V_1.6 channelopathy

As ELIXIR potently inhibited the *I*_NaL_ of the Na_V_1.6 N1768D variant, we next considered whether ELIXIR was able to rect ify the alteration of neuronal AP properties associated with this variant^64^. To do so, we tested the effect of ELIXIR in a mouse model of neurodevelopmental disorder expressing the Na_V_1.6 N1768D variant^65^. Previous studies have shown that these mice develop seizures and present with SUDEP^65^, consistent with clinical phenotypes^21^. Moreover, these mice display aberrant neuronal AP spike properties, including AP prolongation, spike frequency adaptation, and depolarization block, reflecting an increased *I*_NaL_ in various neuronal subtypes^63,64,66,67^. For this, we packaged ELIXIR, and an mCherry expression reporter, into an AAV9 vector and used intracerebroventricular injection to deliver this AAV to cortical neurons of P0 N1768D homozygous knockin (*SCN8A*^N1768D/N1768D^) and wild-type littermate control *(SCN8A*^+/+^) mice. We then targeted layer V pyramidal neurons for whole cell patch clamp electrophysiology recordings in brain slices. Fluorescence imaging confirmed expression of ELIXIR (**Figure 5A**). Comparison of wild-type and N1768D neurons confirmed a prolongation of the APD (**Figure 5B, S8A-S8B**), due to slowing of the repolarizing phase (**Figure S8C-S8D**)^64^. ELIXIR positive neurons in N1768D slices exhibited a significantly reduced APD compared to ELIXIR negative N1768D neurons (**Figure 5B-C; S8B**). The APs of wild-type neurons were found to be insensitive to ELIXIR (**Figure 5B-C; S8B**). Analysis of AP properties showed a small change in AP firing rate in neurons from wild-type mice upon ELIXIR expression (**Figure 5D,5F, S8E**). By contrast, neurons from N1768D mice displayed premature depolarization block, which may contribute to disease phenotypes in *SCN8A* neurodevelopmental disorders^67^, that was significantly attenuated by ELIXIR expression (**Figure 5E, 5G, S8E-S8F**). These results reveal that expression of ELIXIR is effective at reversing the altered neuronal AP properties observed in Na_V_1.6 GoF channelopathies linked to epilepsy.

**Figure 5.**
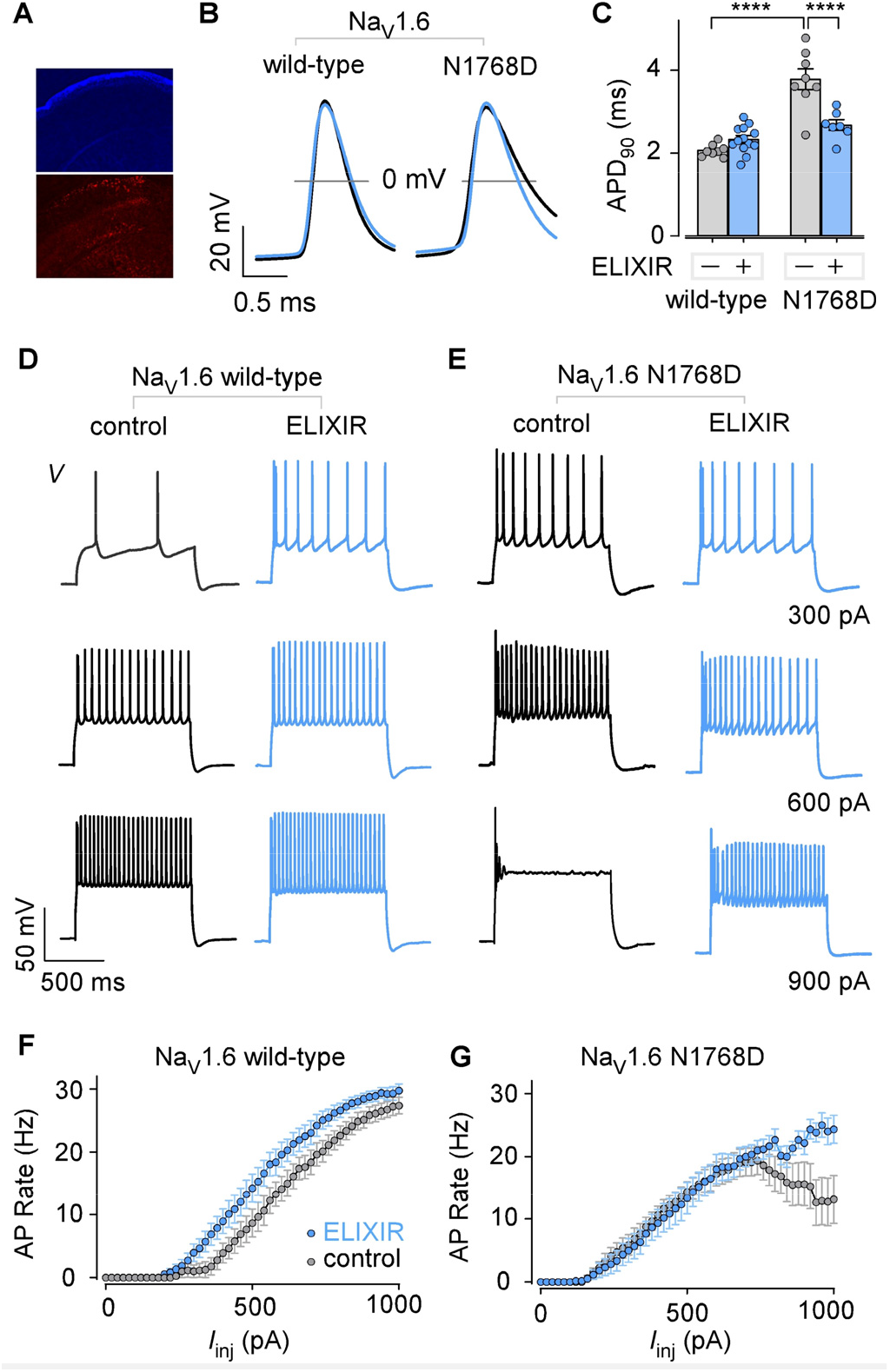
ELIXIR rescues Na_V_ channelopathy-induced AP abnormalities in mouse brain. **(A)** DAPI stain (**top**) and mCherry expression (**bottom**) indicate AAV9-medited expression of ELIXIR in neurons of the cerebral cortex, marked with mCherry fluorescence. **(B)** Group average of AP recordings from control (black trace) and ELIXIR expressing (blue) neurons from WT Na_V_1.6 (**left**) and Na_V_1.6 N1768D (**right**) mice. **(C)** Bar graph summarizing the effect of ELIXIR on the APD_90_ of control and ELIXIR expressing WT Na_V_1.6 (*SCN8A*^+/+^; control: n = 8, N = 2; ELIXIR: n = 8, N = 2) or Na_V_1.6 N1768D (*SCN8A*^N1768D/N1768D^, control: n = 8, N = 2; ELIXIR: n = 7, N = 2) neurons. ****p<0.0001 by two-way ANOVA followed by Tukey’s multiple comparisons test. **(D-E)** Exemplar recordings of APs elicited from either WT (**panel D**) or N1768D (**panel E**) neurons with or without ELIXIR in response to 300 pA (**top**), 600pA (**middle**), and 900 pA (**bottom**) current injection. **(F-G)** Input-output curves depict the AP rate of WT neurons (**panel F**; control: n = 8, N = 2; ELIXIR: n = 8, N = 2) and N1768D neurons (**panel G**; control: n = 8, N = 2; ELIXIR: n = 6, N = 2) in the presence or absence of ELIXIR as a function of applied current. **(K)** Bar graph summarizes effect of ELIXIR on wild-type and mutant Na_V_1.6 channels. Each bar, mean ± SEM. Na_V_1.6 without (n = 10; 973 sweeps) and with ELIXIR (n = 9; 885 sweeps); Na_V_1.6 N1768D mutant without (n = 7; 876 sweeps) and with ELIXIR (n = 10; 1162 sweeps). §*p* = 0.0009 (wild-type Na_V_1.6 and Na_V_1.6 N1768D),, \**p* = 0.0008 (Na_V_1.6 N1768D mutant with and without ELIXIR) by a one-way ANOVA followed by a Tukey’s multiple comparisons test.

### ELIXIR reduces *I*_NaL_ and APD_90_ of CaMKII*δ*_C,OE_ ventricular myocytes

In HF, phosphorylation of Na_V_1.5 by CaMKII results in elevated *I*_NaL_, which contributes to increased risk of cardiac arrhythmias^51,52,68^. As such, we examined whether ELIXIR may reverse pathological *I*_NaL_ in adult mouse ventricular myocytes (aMVMs) isolated from a transgenic mouse model that overexpress the predominant cardiac CaMKII isoform (CaMKII*δ*_C,OE_)^69^. Previous studies have shown that overexpression of CaMKII*δ*_C_ sufficed to induce cardiac hypertrophy and dilated cardiomyopathy leading to arrhythmogenic HF and premature death in these mice^69,70^. Indeed, we observed morphological and electrophysiological changes in the hearts and cardiomyocytes of the CaMKII*δ*_C,OE_ mice (**Figure S9A – S9F**) that were indicative of hypertrophy, and consistent with HF. We utilized ELIXR-cpp for short-term intracellular delivery of ELIXIR into aMVMs (**Figure 6A**, ELIXIR-cpp). Flow cytometric analysis of non-transgenic (nTG) aMVMs treated with ELIXIR-cpp showed a concentration-dependent increase in FITC fluorescence intensity (**Figure 6B-C**), confirming uptake of ELIXIR-cpp by the myocytes. As in previous studies, *I*_NaL_ was quantified through whole-cell patch-clamp recordings as the magnitude of Na^+^ current following 500 ms of depolarization normalized to the whole-cell capacitance. In line with previous studies^68^, the CaMKII*δ*_C,OE_ aMVMs displayed a 2-fold increase in *I*_NaL_ relative to aMVMs from nTG animals (**Figure 6D-F**). Application of ELIXIR-cpp caused a small but statistically significant reduction in the *I*_NaL_ of nTG myocytes (**Figure 6D-F**). By comparison, incubation with ELIXIR-cpp had a much stronger effect on the *I*_NaL_ of CaMKII*δ*_C,OE_ aMVMs, which was reduced to near wild-type levels (**Figure 6E-F**).

**Figure 6.**
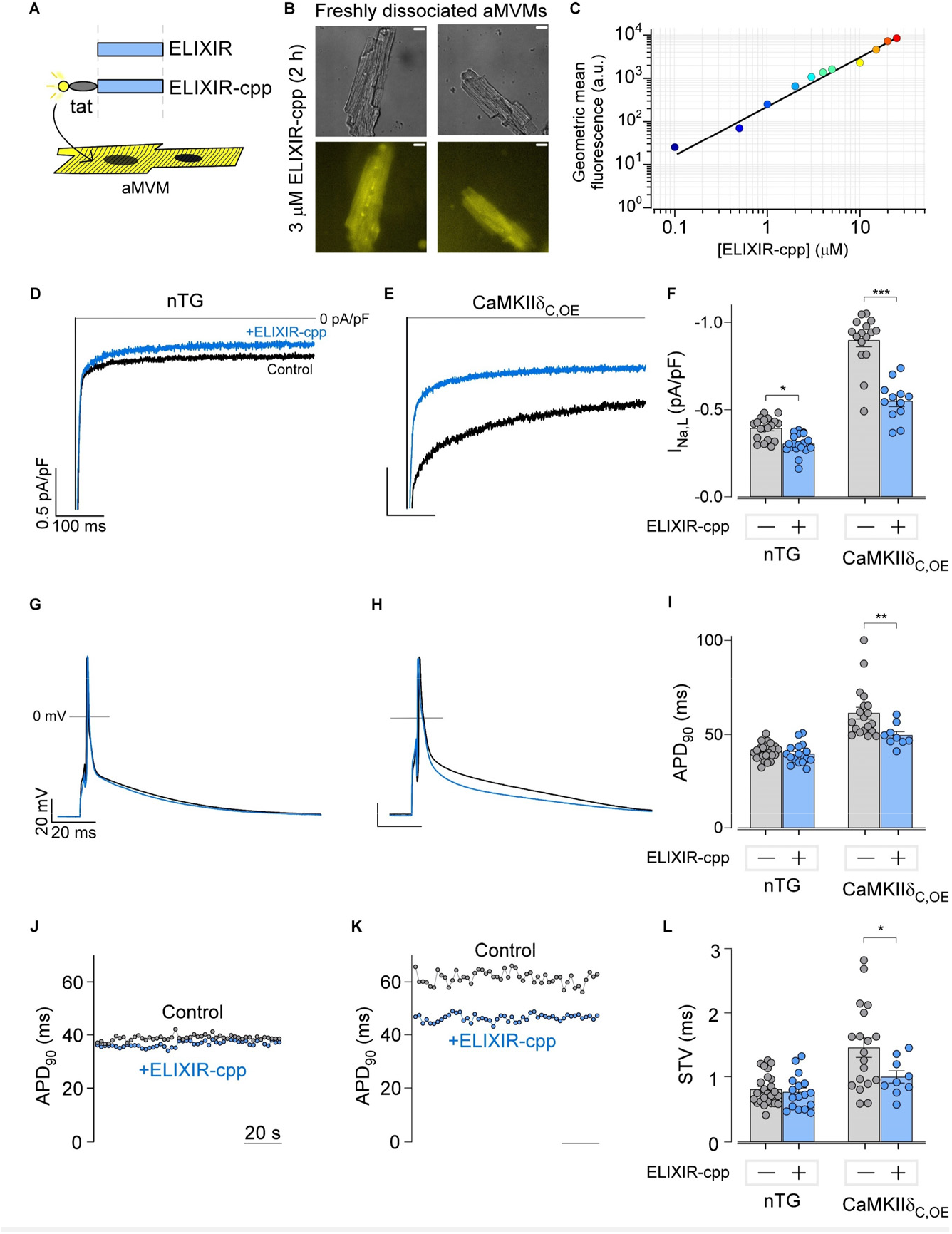
ELIXIR inhibits I_NaL_ and reduces APD_90_ in murine ventricular cardiomyocytes. **(A)** Schematic shows design of cell penetrating ELIXIR (ELIXIR-cpp). We attached a viral tat sequence to the amino terminus of ELIXIR as well as a FITC fluorophore to monitor cellular uptake. **(B)** Epifluorescence images show uptake of ELIXIR-cpp into freshly dissociated ventricular myocytes from adult mice. Scale bar, 1 μm. **(C)** Flow cytometric analysis shows concentration dependent uptake of ELIXIR-cpp into aMVMs. **(D)** Exemplar whole cell recordings show *I*_NaL_ in freshly dissociated aMVM from non-transgenic myocytes (nTG) at baseline (black trace) and following treatment with 3 μM ELIXIR-cpp (blue trace). **(E)** Exemplar whole cell recordings show *I*_NaL_ inhibition in aMVM from a CaMKII*δ*_C,OE_ transgenic mice that develops heart failure. Format as in panel **D. (F)** Bar graph summarizes the effect of ELIXIR on *I*_NaL_ in MVM from nTG versus CaMKII*δ*_C,OE_ mice. Each bar, mean ± SEM. nTG without (n = 19 from 6 mice) or with ELIXIR-cpp (n = 19 from 6 mice); CaMKII*δ*_C,OE_ without (n = 16 from 5 mice) or with ELIXIR-cpp (n = 12 from N = 5 mice). ^*^*p* = 0.034, \*\*\**p* < 0.001 by a two-way ANOVA followed by a Tukey’s multiple comparisons test. **(G – H)** Exemplar AP recordings from nTG (**G**) and CaMKII*δ*_C,OE_ (**H**) aMVMs. Black trace, without ELIXIR; Blue trace, with ELIXIR. **(I)** Bar graph summarizes ELIXIR effect on APD_90_. Each bar, mean ± SEM. nTG without (n = 25 from 6 mice) or with ELIXIR-cpp (n = 17 from 6 mice) and CaMKII*δ*_C,OE_ without (n = 19 from 5 mice) or with ELIXIR-cpp (n = 9 from 4 mice) \*\**p* = 0.004 by two-way ANOVA followed by Tukey’s multiple comparisons test. **(J – K)** Diary plot shows variability in action potential durations (APD90) for non-transgenic or CaMKII*δ*_C,OE_ transgenic mice in the presence and absence of ELIXIR. **(L)** Bar graph summarizes changes in short term variability (STV) of APDs.

To further probe the potential functional consequences of ELIXIR-cpp on cardiomyocyte function, we undertook current clamp recordings to quantify changes in AP morphology. Consistent with the increase in *I*_NaL_, the AP duration (APD) of CaMKII*δ*_C,OE_ myocytes was significantly prolonged relative to that of nTG myocytes (**Figure 6G–I**). In addition to this, myocytes from the CaMKII*δ*_C_,_OE_ mice had greater variability in the APD, quantified as short-term variability^71,72^ (STV, **Figure 6J–L**), than those from nTG animals consistent. Remarkably, application of ELIXIR-cpp yielded a significant reduction in the APD (**Figure 6H-I**) and STV (**Figure 6K-L**) of aMVMs from CaMKIId_C_,_OE_ mice, reversing both to near nTG levels. By comparison, application of ELIXIR-cpp caused essentially no change in APD or STV of nTG aMVMs. Collectively, these findings illustrate the ability of ELIXIR, applied extracellularly as a peptide, to tune native channel function.

### ELIXIR Inhibits *I*_NaL_ in LQT3 patient derived iPSC-CMs

While ELIXIR-cpp allows for short-term modification of Na_V_ function in cardiomyocytes, constitutive expression of ELIXIR may be advantageous in pathophysiological settings. As such we used AAV9 to express ELIXIR in two cardiac disease models: (1) human cardiomyocytes differentiated from induced pluripotent stem cells (hiPSC-CMs) and (2) transgenic mice expressing a mutant Na_V_1.5 (F1759A^tg^) (**Figure 7A**).

**Figure 7.**
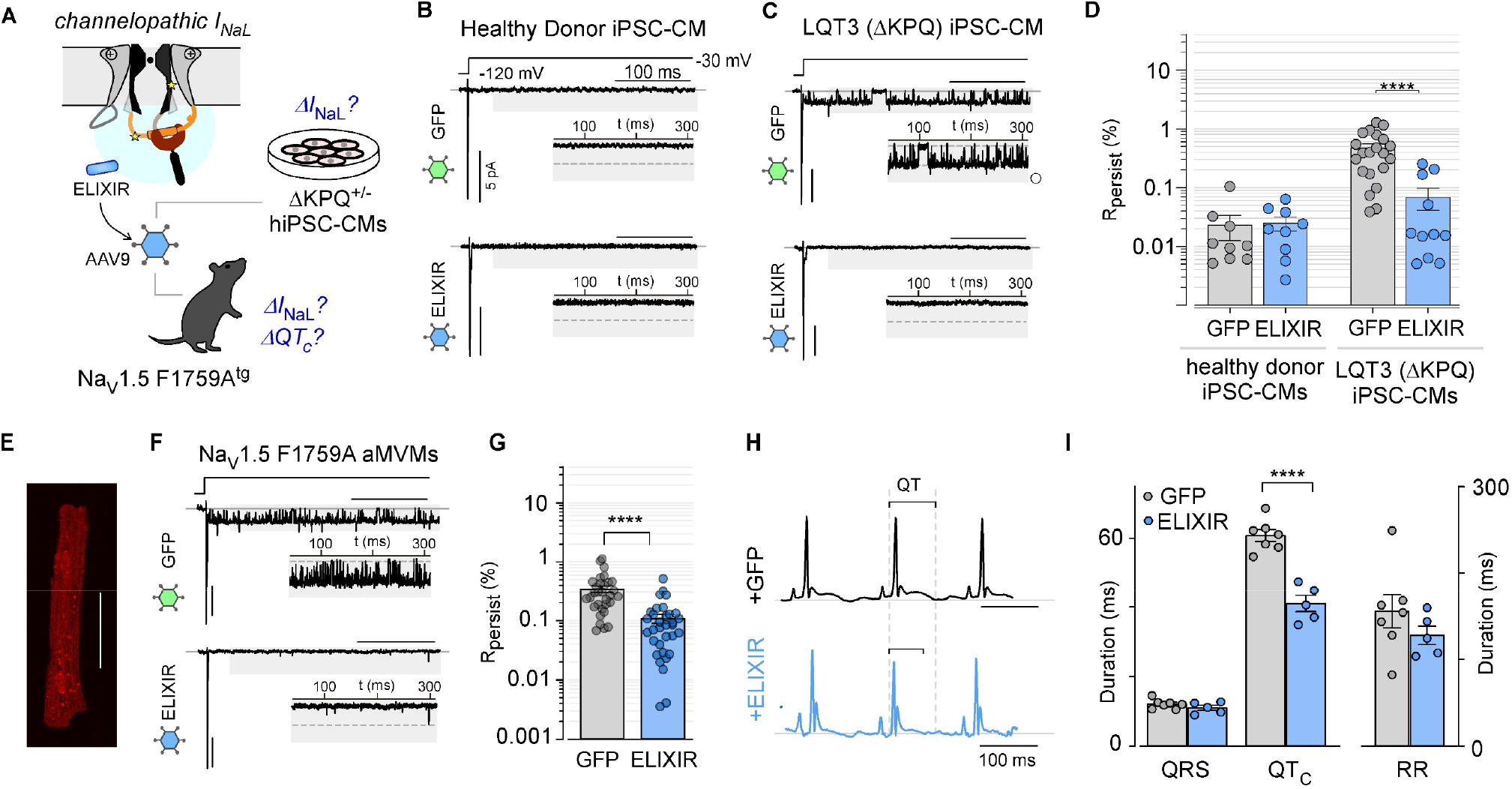
ELIXIR reverses Na_V_1.5 dysfunction in LQT3 hiPSC-CMs and in F1759A^tg^ mice. **(A)** Schematic shows AAV9 mediated expression of ELIXIR into cultured LQT3 hiPSC-CMs and F1759A^tg^ mice to determine changes in *I*_NaL_ or electrocardiogram. **(B - C)** Exemplar multichannel recordings of iPSC-CMs differentiated from a healthy donor (**A**), or from an LQT3 patient carrying a heterozygous ΔKPQ mutation (**B**) transduced with an adenovirus containing GFP (top) or ELIXIR (bottom). Insets show channel openings in the late phase enlarged for better visualization. **(D)** Bar graph summarizes effect of ELIXIR on *I*_NaL_ in iPSC-CMs. Each bar, mean ± SEM. HD iPSC with GFP (n = 9; 781 sweeps from N = 2 differentiations) or ELIXIR (n = 9; 803 sweeps from N = 2 differentiations); LQT3(ΔKPQ) iPSC-CMs with GFP (n = 21; 1361 sweeps from N = 2 differentiations) or ELIXIR (n = 11; 885 sweeps from N = 2 differentiations). \*\*\*\**p*<0.0001 by a two-tailed Mann-Whitney U-test. **(E)** Confocal image shows mCherry expression as a marker of ELIXIR production by ventricular myocytes freshly isolated from adult Na_V_1.5 F1759A mice. Scale bar, 50 μm. **(F)** Exemplar multichannel recordings from ventricular myocytes isolated from Na_V_1.5 F1759A mice infected with an AAV containing GFP (top) or ELIXIR (bottom). Insets show channel openings in the late phase enlarged for better visualization. **(G)** Bar graph summarizes effect of ELIXIR on *I*_NaL_ of Na_V_1.5 F1759A aMVMs. Each bar, mean ± SEM. Na_V_1.5 F1759A mice injected with GFP AAV (n = 31; 2,181 sweeps from N = 6 mice) or ELIXIR AAV (n = 33; 2,396 sweeps from N = 6 mice). \*\*\*\**p*<0.0001 by a two-tailed Mann-Whitney U-test. **(H)** Exemplar limb-lead surface ECGs of isoflurane-anesthetized F1759A mice infected with an AAV containing GFP (top) or ELIXIR (bottom). **(I)** Bar graph comparison of the RR (**H**), and QRS and QT_C_ (**I**) intervals of isoflurane-anesthetized F1759A mice infected with an AAV containing GFP (N = 7 mice) or ELIXIR (N = 5 mice). \*\*\*\**p*<0.0001 by nested t-test.

First, we considered hiPSC-CMs derived from an LQT3 patient heterozygous for the Na_V_1.5 ΔKPQ variant^26^, as well as those from a healthy donor (HD)^73^. Following differentiation and maturation, spontaneous contractions were observed in both the HD and ΔKPQ hiPSC-CMs. Multichannel recordings of the HD hiPSC-CMs revealed low baseline levels of *I*_NaL_, that were minimally altered by AAV9-mediated expression of ELIXIR (**Figure 7B, S9G**). By contrast, the ΔKPQ hiPSC-CMs exhibited markedly increased (∼60-fold) *I*_NaL_ (**Figure 7C-D, S9H**). Following transduction with ELIXIR, a ∼7-fold reduction in *I*_NaL_ was observed in the ΔKPQ hiPSC-CM (**Figure 7C-D, S9H**). This underscores the efficacy of ELIXIR in diminishing *I*_NaL_ linked to LQT3 variants in the more pathophysiologically relevant setting of human cells.

Second, we considered a transgenic mouse model that inducibly expresses the lidocaine-resistant Na_V_1.5 F1759A mutant channel. These mice have been previously reported to have an elevated *I*_NaL_ and prolonged QT interval^2^. Subcutaneous injection delivered an AAV9 encoding ELIXIR or GFP to P4 mice. Eight weeks after the injection of the virus, ventricular cardiomyocytes were isolated, and ELIXIR expression was confirmed by confocal imaging of mCherry fluorescence (**Figure 7E**). Multichannel recordings of cardiomyocytes from GFP-injected mice revealed the presence of late channel openings (**Figure 7F**), with an *I*_NaL_ consistent with previous reports^25^. By comparison, cardiomyocytes from ELIXIR-injected animals showed a significantly attenuated *I*_NaL_ compared to those from GFP control mice (**Figure 7F-7G**). As increased *I*_NaL_ from Na_V_1.5 manifests as a prolonged QT interval on the ECG and is the basis for LQT3 in humans, we obtained limb-lead surface electrocardiographic (ECG) recordings from both ELIXIR injected mice and GFP-control littermates (**Figure 7H**). QT intervals, corrected for heart rate using Mitchell’s formula (QT_C_)^74^, were significantly shorter in the ELIXIR expressing mice than in the GFP-expressing control animals (**Figure 7I**). We observed no significant differences in the RR or QRS intervals (**Figure 7I**). Collectiv ely, these results confirm the effectiveness of long-term ELIXIR expression in tuning endogenous *I*_NaL_ in both human and murine models of Na_V_1.5 dysfunction.

## Discussion

Strategies to precisely manipulate ion channel function are highly sought after both as tools for unraveling their physiological functions and for developing new therapies^75^. A prominent example of this is the family of Na_V_ channels (Na_V_1.1-1.9), where gain-of-function variants have been associated with a broad range of human diseases including cardiac arrhythmia^76^, myotonia^6^, and epilepsy^21^. Despite their varied clinical presentations, many of these diseases stem from a common deficit in Na_V_ channel inactivation that leads to sustained Na^+^ influx. By combining structural insights and a computational protein design framework, we developed a peptide modulator of Na_V_ channels designed to promote inactivation, called ELIXIR. Our biochemical and electrophysiological analysis showed that ELIXIR binds to the Na_V_1.5_CTD_ with submicromolar affinity and can modulate the “pathological” *I*_NaL_ of multiple Na_V_ isoforms. We further demonstrate the efficacy of this synthetic *I*_NaL_ modulator to reverse pathophysiological changes in multiple disease models linked to Na_V_ channel dysfunction. These findings suggest that ELIXIR could serve as a template for developing a new class of rationally designed Na_V_ channel modulators and further highlight the potential utility of *de novo* protein design for custom engineering of ion channel modulators in health and disease.

Many ion channels are large membrane-spanning multi-subunit complexes, the gating of which is controlled by intricate conformational dynamics. Consequently, engineering of synthetic ion channel modulators with desired functional effects has been challenging. To date, the prevailing strategy has been largely empirical, and focused on repurposing known ion channel modulators^26,77^ or developing synthetic nanobodies that interact with key channel domains^78^. Recently, major advances in computational protein design, particularly deep learning approaches, have improved accuracy while also reducing the required computational time^27-29,43,79^, suggesting that these methods may provide alternate avenues for developing novel ion channel modulators. Furthermore, the availability of open-source and easy-to-use platforms has increased the accessibility of these methods to the broader scientific community. While promising, experimental validation of *de novo* sequences generated by these algorithms and establishing their functionality in native biological settings has been challenging. For in silico designed *de novo* protein-binders, only a small number of targets have been tested *in vitro*^28,29^ and even fewer *in situ* or *in vivo*^80^. Here, we focused on Na_V_1.5 channels owing to their high pathophysiological relevance. We targeted the Na_V_1.5_CTD_, as this region has emerged as a regulatory hub that interacts with multiple auxiliary proteins to fine tune channel function. Furthermore, the Na_V_1.5_CTD_ also contains a large hydrophobic pocket that provides an attractive interface for binder design. With the use of the design protocol described here, we were able to generate a *de novo* peptide sequence that binds the Na_V_1.5_CTD_ and tunes a specific aspect of Na_V_1.5 function (i.e. *I*_NaL_) in heterologous expression systems, primary cardiomyocytes and *in vivo*. Our results highlight the applicability of these methods for developing novel ion channel modulators and lend further support to the general utility of currently available algorithms for targeting a wide array of proteins.

In the heart, multiple pathophysiological mechanisms have been linked to increased *I*_NaL_. These include: (1) human Na_V_1.5 variants distributed across various channel domains^81^, (2) channel phosphorylation by CaMKII or PKA^51,52^, (3) reactive oxygen species (ROS)^52^ or cellular metabolites^82^, (4) altered binding of channel interacting proteins^48,83-85^, and (5) diverse cell signaling mechanisms^86-88^. As such, considerable effort has been devoted to developing selective *I*_NaL_ inhibitors. In this regard, class 1b antiarrhythmics (e.g. lidocaine and mexiletine), which bind with rapid kinetics to a transmembrane site in Na_V_1.5, exhibit some preference for inhibiting *I*_NaL_^9^. Ranolazine, structurally similar to lidocaine, has been found to exhibit an even greater preference for inhibiting *I*_NaL_. High-resolution structures have been solved of Na_V_1.5 bound to several of these antiarrhythmics^23,24^, and in each, the drug is bound to a nearly identical site in the channel pore. Importantly, this shared binding site likely limits the selectivity of these molecules for late versus peak Na^+^ current as their interaction obstructs ion flux through the pore. Instead of targeting the pore domain, our approach was to leverage the natural inactivation mechanism of Na_V_ channels. Specifically, we sought to promote inactivation by enhancing the release of the IG from the Na_V_1.5 CTD through the application of a competitive binder to this site (ELIXIR). This strategy has two advantages. First, by not targeting the channel pore or the voltage-sensing domains, ELIXIR is unlikely to significantly perturb channel activation. Experimentally, we found that ELIXIR had minimal effect on peak current density and voltage-dependence of activation (**Figure S3D – S3F**). Second, ELIXIR is unlikely to exert substantial effects on channels that have no baseline defect in inactivation. Specifically, for wild-type channels, the DIV VSD movement is considered a rate-limiting step for both the development of and recovery from inactivation^34^. Without the movement of DIV VSD, it is unlikely that the pore-proximal site is available for interaction with the IFM motif. In this case, displacement of the IG by ELIXIR may not promote inactivation. One interesting possibility hinted at by our results is that in pathophysiological settings, the release of IG from the CTD may become a limiting step in addition to DIV VSD movement. As a result, promoting IG release by ELIXIR would be expected to boost inactivation. This aspect is potentially important as it could lend ELIXIR selectivity for tuning pathogenic *I*_NaL_. Experimentally, we found that ELIXIR had minimal effects on *I*_NaL_ of unmodified wild-type Na_V_. channels. Nevertheless, ELIXIR was still effective at inhibiting *I*_NaL_ resulting from (1) diverse Na_V_1.X mutations (**Figure 3A – 3E, 4H, 4K**) and (2) channel phosphorylation by PKA or CaMKII (**Figure 3D-E**). A corollary is that the generality of ELIXIR activity on Na_V_s suggests that these seemingly distinct perturbations all converge on a common endpoint, the diminished ability of the IG to bind the pore-proximal site. For some Na_V_1.5 perturbations examined, ELIXIR did not fully reverse the increase in *I*_NaL_. This partial inhibition could stem from incomplete disruption of the IG-CTD interaction by ELIXIR or may reflect other structural defects caused by these perturbations. Further optimization of ELIXIR through affinity maturation or the production of mutation-specific binders could help address these limitations. Of note, we recently found that the endogenous Na_V_ channel interacting protein Fibroblast growth factor homologous factors (FHFs) inhibits *I*_NaL_ in an isoform/splice-variant specific manner^26^. Based on this, we engineered a 39-amino acid FHF1_A_ fragment (FHF-inhibiting X-region, FixR) as an *I*_NaL_ inhibitor. While the molecular mechanism by which FHFs/FixR reduce *I*_NaL_ remains unclear, it appears likely that it is distinct from that of ELIXIR, as FHFs can promote inactivation of both the Na_V_1.5 Δ1810 variant that lacks the ELIXIR-binding pocket and mutations that disrupt IG (IFM/IQM) ^26^. As such, it is possible that the two peptides may be beneficial in different settings depending on the specific perturbations in channel function. Also, as FHFs are natively present in various tissues, the effectiveness of FixR may vary depending on the level of FHF expression. By contrast, ELIXIR as a de novo peptide could be more broadly applicable.

Beyond cardiac maladies, Na_V_ channel malfunction has been implicated in a variety of diseases including epilepsy syndromes^64^, myotonia^6^, and chronic pain^22^. The high degree of sequence similarity in the Na_V_ EFL suggests that ELIXIR may function as an *I*_NaL_ inhibitor for channelopathies across the Na_V_ family, an attractive possibility given the range of diseases associated with dysfunction of these channels (**Figure 4A-B**). Supporting this idea, we found that co-expression of ELIXIR was sufficient to reverse the increase in *I*_*NaL*_ observed with both the Na_V_1.4 F1698I variant linked to a cold-aggravated myotonia as well as the well-studied Na_V_1.6 N1768D variant linked to epilepsy (**Figure 4F – 4K**). Consistent with previous studies^21,63,64^, slice electrophysiological recordings layer V cortical neurons from homozygous N1768D mice showed changes in key AP properties of including slower repolarization of the AP and increased presence of depolarization block (**Figure 5**). Viral expression of ELIXIR was sufficient to partially reverse these changes in neuronal AP morphology and excitability, pointing to the potential effectiveness of ELIXIR in the nervous system. More broadly, the Na_V_ CTD has an architecture similar to the C-termini of other ion channels, including the voltage-gated calcium channels^15^ and the sodium leak channel NALCN^14^. In high-resolution structures of these channels, the EFL region of the CTD is bound to the domain III – domain IV linker^14,15^, hinting at conserved modulatory mechanisms. If so, our design strategy may be broadly effective for developing selective actuators of this physiologically important channel superfamily.

Three key limitations merit further attention. First, a high-resolution structure of ELIXIR bound to an Na_V_ channel is not available. Nevertheless, structural predictions of ELIXIR/CTD complexes by various algorithms all converge to a nearly identical interface. Experimentally, the mutational analysis of ELIXIR residues was consistent with the predicted interface. Second, although peptides are used clinically (e.g. semaglutide and insulin), the specific translational application of ELIXIR may be impaired by the requirement for intracellular delivery. Our studies utilized two possible approaches, namely fusion to a cell-penetrating peptide and viral delivery of ELIXIR. Both approaches were effective for robust delivery and reversal of *I*_NaL_ in different settings. Nevertheless, these strategies have potential limitations for translational applications including immunogenicity. Other emerging protein delivery approaches may also be beneficial for ELIXIR. Future studies will address these possibilities and assess the effectiveness of potential intracellular delivery mechanisms. Alternatively, it may be possible to identify small molecules that interact with the IG-binding pocket of an Na_V_1.x CTD and mimic the action of ELIXIR. In this regard, the change in fluorescence anisotropy with ELIXIR may be advantageous for establishing in vitro high-throughput screens. Third, with viral expression of ELIXIR, we were unable to estimate a concentration as the mCherry marker is bicistronically expressed. It is possible that the concentration of ELIXIR may be supramaximal, which may cause off-target effects. Future *in vivo* studies will consider the dose-dependence of ELIXIR expression.

Although ion channel dysfunction has been long associated with various life-threatening diseases, these proteins have often been considered challenging targets for drug development^89^. A key difficulty is that in pathophysiology, the functional change often involves alterations in dynamic channel behavior that disrupts cellular excitability. As such, simply turning an ion channel on or off is often insufficient. Our results demonstrate the effectiveness of computational protein design for the development of synthetic proteins/peptides that tune channel dynamics. This ability could turbocharge the creation of innovative strategies to directly target the pathophysiology of multiple channelopathies and enable precision engineering of nature’s transistors.

## Materials and Methods

### Molecular biology

The human Na_V_1.5 sequence used in this work corresponds to GenBank accession number M77235.1 and was expressed from a pGW vector, as described previously^26^. The IQ/AA mutation in the channel CTD and ΔKPQ mutation in the inactivation gate were generated using overlap-extension PCR followed by ligation into the NruI and XbaI, and KpnI and XbaI restriction sites, respectively. The Δ1864 truncation was made by PCR amplifying the relevant segment using a forward primer upstream of the KpnI site and reverse primer that truncated at position 1864, which included an XbaI site for ligation into the Na_V_1.5/pGW construct. A gene fragment containing the DNA sequence of the designed peptide (MSPRREALYRGFRACYDVLRH) was purchased from Synbio technologies and subcloned into a pIRES2-eGFP vector using the NheI and BamHI restriction sites. For FRET experiments the Venus-tagged Na_V_1.5_CTD_ (Na_V_1.5 residues 1771-2016) and Cerulean-tagged ELIXIR (WT, L8A, F12E, and Y16A) were expressed from a pCDNA3 vector. The PKA catalytic domain and CaMKII_T286D_ were expressed from a pCDNA3.1 plasmid as previously reported^90^. The WT Na_V_1.1, Na_V_1.4, Na_V_1.6, and Na_V_1.7 were expressed from pIR-CMV-IRES-mScarlet, pCDNA3.1, pCDNA4TO-IRES-mScarlet, and pCMV plasmids, respectively. For bacterial overexpression and purification, the Na_V_1.5_CTD_ (Na_V_1.5 residues 1773-1940) was expressed as a GST-fusion inserted between the BamHI and XhoI sites of a pGEX-6P vector. Calmodulin was cloned into a pRSF plasmid using the NcoI and XhoI sites. QuickChange mutagenesis was used to substitute the single cystine residue (C15) with a serine to generate the final ELIXIR peptide sequence (MSPRREALYRGFRASYDVLRH), to produce the binding-deficient ELIXIR mutants (L8A, F12E, Y16A) and the Na_V_1.6 N1768D mutant channel.

### Peptide design

Peptide design was done using the AfDesign algorithm implemented in ColabDesign^31^. The coordinates for the Na_V_1.5_CTD_ were taken from an x-ray crystallographic of the Na_V_1.5_CTD_ bound to calmodulin and FHF2_U_ (PDB ID: 4DCK)^44^. For the first round of design, residues 1786 - 1802 and 1850 - 1887 of Na_V_1.5 were used as a hotspot, the binder length was set to 21 residues, and the number recycles, and models used were 1 and 2, respectively. The design protocol was then run with a semi-greedy optimizer (pssm_semigreedy) and the default gradient decent settings. The pssm_semigreedy protocol starts by optimizing, via gradient descent, a softmax representation of the sequence. This is done for 120 steps. This results in a vector of shape binder length (21) that we call PSSM (position specific scoring matrix), which represents the probability of each amino acid at each position. For the starting sequence, the amino acid at each position is selected that maximizes the probability. For the semi-greedy protocol, at each step, 10 independent mutations are evaluated and the mutation with best loss is fixed. The positions are sampled following 1-pLDDT^91^ and the amino acid identity following PSSM. This is done for 32 steps. The sequence with best loss across the 32 steps is selected. The loss combines pLDDT over the binder positions plus contact score: 0.1 * (1-pLDDT) + 1.0 * contact score. The contact score minimizes the entropy of the distogram output (distance distribution between c*β* atoms) between the target and the peptide. More specifically, every peptide position must have at least one confident distance prediction to any target position, unless a hotspot is defined. The protocol is available via Google Colab at: https://colab.research.google.com/github/sokrypton/ColabDesign/blob/v1.1.1/af/examples/peptide_binder_design.ipynb. Following the first round of design, the final peptide sequence was used as a seed to initialize the next round. This process was repeated a total of five times. The peptide sequence was then screened computationally by comparing the interface between the Na_V_1.5_CTD_ and the peptide in models of the complex generated with ColabFold^41^, ESMFold^42^, OmegaFold^27^ and RoseTTAFold^43^. Contacts of structural units analysis was done with the COCOMAPS webserver^92^ using cutoffs of 6.0 Å and 4.5 Å.

### Protein expression and purification

For protein expression *Escherichia Coli* BL21 (DE+) cells were co-transformed with plasmids for both the Na_V_1.5_CTD_ and CaM and grown in 0.5 L cultures of CIRCLEGROW supplemented with carbenicillin (0.1 mg/mL) and kanamycin (0.05 mg/mL). The cells were grown at 37 °C until OD_600_ of 0.7 was reached at which point expression was induced by the addition of 0.5 mL of 1 M IPTG and the temperature was set to 18°C. Cells were harvested the following morning by centrifugation (4,000 rpm, 4°C, 15 min) and stored at -20°C.

The Na_V_1.5_CTD_ was purified as described previously^93^. Cell pellets were thawed and resuspended in lysis buffer (50 mM HEPES, 500 mM KCl, 50 μM EGTA, 0.01% (w/v) NaN_3_, pH 7.4), disrupted via emulsification, and centrifuged (25,000 rpm, 4 °C, 50 min). The supernatant was then filtered (0.2 μM), mixed with 5 mL of Glutathione Sepharose 4B, and rocked at 22°C for 30 min. The resin was then rinsed with 100 mL of lysis buffer, and the Na_V_1.5_CTD_-GST+CaM complex was eluted in 15 mL of elution buffer (lysis buffer + 10 mM glutathione). The GST-tag and glutathione were removed by the addition of precision protease and overnight dialysis against lysis buffer, respectively. The sample was then mixed with 5 mL of glutathione resin and the Na_V_1.5_CTD_+CaM complex was collected and concentrated. Anion exchange chromatography (pH 8.5 to 7.4 and KCl 0 to 250 mM) was then used to separate the Na_V_1.5_CTD_ and CaM. Protein purity (≥95%) was assessed by SDS-PAGE and UV-vis spectroscopy. Protein concentration was determined from the absorbance at 280 nm.

### Fluorescence anisotropy

The ELIXIR peptide, with an N-terminal fluorescein and TAT sequence (YGRKKRRQRRRGSGMSPRREALYRGFRASYDVLRH, theoretical MW: 4826.48 g/mol), was synthesized by GenScript. Binding of the Na_V_1.5_CTD_ to the ELIXIR peptide (100 nM) was monitored in 50 mM HEPES, 100 mM KCl, 1 mM MgCl_2_, 100 mM EGTA, 0.01% (w/v) NaN_3_, 1 mM DTT, pH 7.4 at 22 °C. The fluorescence anisotropy (*λ*_em_ = 494 nm, *λ*_ex_ = 518 nm, 5 nm excitation and emission bandpasses, 4 sec integration) was monitored throughout the titration with a fluorolog fluorometer (Horiba). Anisotropy (R) was calculated with,

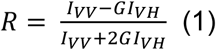

where I_VV_ and I_VH_ are the intensities of the vertically excited and vertically emitted, and vertically excited and horizontally emitted light, respectively. G was determined prior to each titration as the ratio between the horizontally excited and vertically emitted light, and the horizontally excited and horizontally emitted light (I_HV_/I_HH_). Each data point was the average of the three measurements.

The normalized anisotropy was fit to Eq 2 with the Solver add-in in Excel,

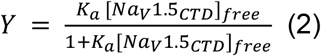

where [Na_V_1.5_CTD_]_free_ was determined with Eq 3

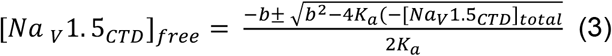

In Eq 3, b was (1+K_a_[ELIXIR peptide]-K_a_[Na_V_1.5_CTD_]_total_). The reported K_D_ and standard deviation were determined from 3 replicate titrations.

### Cell culture and transfection

HEK293 cells (ATCC CRL1573) were cultured in 60-mm dishes on glass coverslips and transfected with the Ca^2+^-phosphate method. For electrophysiology recordings, cells were co-transfected with 4-10 μg of cDNA encoding the desired Na_V_ channel variant, 1 µg of YFP, and 0.5 μg of simian virus 40 T-antigen ± 3 μg of ELIXIR. For experiments investigating the phosphorylation-dependent late Na^+^ current, 3 μg of cDNA encoding the catalytic domain of PKA or CaMKII_T286D_ was used. Four hours after transfection, the cells were rinsed, and the media replaced. Electrophysiological recordings were performed at room temperature 1 – 2 days after transfection.

### Whole-Cell recording from HEK293 cells

Whole-cell electrophysiology recordings were conducted at room temperature with an Axopatch 200B amplifier (Axon Instruments). Pipettes contained (in mM): 114 CsMeSO_3_, 5 CsCl, 1 MgCl_2_, 4 Mg-ATP, 10 HEPES (pH 7.4), and 10 BAPTA (1,2-bis(o-aminophenoxy)ethane-N,N,N′,N′-tetraacetic acid), at adjusted to 290 mOsm with glucose. The bath contained (in mM) 20 NaCl, 1 CaCl_2_, 10 HEPES, pH 7.4 (adjusted with NaOH), at 300 mOsm (adjusted with glucose). Pipettes were pulled from borosilicate glass (World Precision Instruments, MTW 150-F4), with a horizontal micropipette P-97 puller (Sutter Instruments) and fire polished with a microforge (Narishige). Pipettes typically had a resistance of 2 – 3 MΩ, which was >70% compensated. An ITC-18 (Instrutech), controlled by custom MATLAB software (Mathworks), was used during data acquisition. Currents were low-pass filtered at 2 kHz before being digitized at several times that frequency. P/8 leak subtraction was used. Cells were kept at a holding potential of −120 mV.

### Multichannel recordings

Multichannel recording from HEK293 cells, iPSC-CMs, or aMVMs were obtained in the on-cell configuration as in our previous work^25,26^. Pipettes contained a buffer comprised of (in mM): 140 NaCl, 10 HEPES, 1 CaCl_2_, pH 7.4 (adjusted with tetraethylammonium hydroxide), and at 300 mOsm (adjusted with tetraethylammonium methanesulfonate). To zero the membrane potential, the bath contained (in mM): 132 K-glutamate, 5 KCl; 5 NaCl, 3 MgCl, 2 EGTA, 10 glucose, 20 HEPES, pH 7.4 (adjusted with NaOH), at 300 mOsm (adjusted with glucose). For experiments monitoring the dose-dependent effect of ELIXIR on I_NaL_, HEK293 cells were treated with the indicated concentration of ELIXIR-cpp at 30 °C for 4 hours after which multichannel recordings were collected. Recordings were conducted at room temperature using the integrating mode of an Axopatch 200A amplifier (Axon Instruments, Molecular Devices). Patch pipettes were pulled from ultra-thick-walled borosilicate glass (BF200-116-10; Sutter Instruments) using a horizontal puller (P-97, Sutter Instruments), fire polished with a microforge (Narishige), and coated with Sylgard (Dow Corning). Pipettes typically had a 4 – 7 MΩ resistance. Elementary currents were low-pass filtered at 2 kHz with a four-pole Bessel filter and digitized at 200 kHz with an ITC-18 (Instrutech) analog-to-digital converter, controlled by custom MATLAB software (Mathworks). For every pulse P/8 leak pulses was obtained. Leak subtraction was done with an automated algorithm that fit the kinetics of the leak current and capacitive transient using convex optimization with L1 regularization of the following objective function in equation 4:

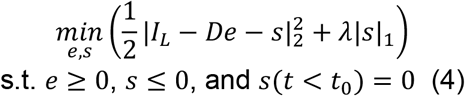

Here, *I*_L_ is the leak waveform that is to be fit as a sum of exponentials; *D* is a matrix composed of a library of exponential functions with various time constants; *e* is a sparse vector composed of the amplitude of each exponential; *s* is a vector representing an estimate of negative outliers from the baseline that correspond to channel openings; and λ is a penalty term set to be 0.25. After leak subtraction, the unitary current for each patch was estimated from an amplitude histogram and each trace was subsequently idealized. The ensemble average from 50 - 120 stochastic traces was computed for each patch and normalized to the peak current. The average late current for each patch (*R*_persist_) was computed as the average normalized *P*_O_ following 50ms of depolarization. Concentration-response data were analyzed using a nonlinear least squares fit with a Hill equation (Graphpad Prism 10). The upper and lower plateau values of the curve were fixed to the R_persit_ value of Na_V_1.5 ΔKPQ expressed alone (upper) and co-expressed with ELIXIR (lower).

### Flow cytometric FRET

The flow cytometric two-hybrid binding assays were performed as previously described^94^. HEK293 cells (ATCC CRL1573) cultured in 12-well plates were transfected with linear polyethylenimine 25 kDa polymer (PEI, Polysciences #2396602) using a 2:1 PEI to DNA mass ratio. For each experiment cells were transfected with 3 μg Venus-Na_V_1.5_CTD_, 1 μg Cerulean-ELIXIR, and 0.5 μg T antigen, after which cells were cultured for 2 days. To enhance the level of mature fluorophores protein production was inhibited with Cycloheximide, 100 μM, 2 hours before fluorescence measurements were made.

Fluorescence was measured with an LSRII (BD Biosciences) flow cytometer, equipped with 405 nm, 488 nm and 633 nm lasers for excitation and 18 emission channels. Healthy single cells were selected based off the forward and side scatter signals. FRET efficiency was determined by measuring three different fluorescent signals: 1) Cerulean emission following direct excitation (S_Cer_) through the BV421 channel (excitation 405 nm, emission 450/50), 2) Venus emission through direct excitation (S_Ven_) with the fluorescein isocyanate channel (excitation 405 nm, dichroic 505 LP, emission 525/50) and 3) Venus emission due to FRET (S_FRET_) through the BV510 channel (excitation 405 nm, dichroic 505 LP, emission 525/50). These measurements were used to obtain Cer_direct_ (Cerulean emission due to direct excitation), Ven_direct_ (Venus emission due to direct excitation) and FRET efficiency. Flow cytometric signals were collected at a medium rate (2,000 to 8,000 events/sec) and the data were exported as Flow Cytometry Standard 3.0 files for further processing and analysis with custom MATLAB software (Mathworks).

As previously described^94^, several control experiments were conducted for each experimental run of the flow cytometer. Firstly, the background fluorescence level of each channel (BG_Cer_, BG_Ven_, and BG_FRET_) was determined using cells that were not exposed to any fluorophore containing plasmid. Secondly, cells expressing only Venus were used to determine the spectral crosstalk parameter *R*_*A1*_, which corresponds to the bleed-through of Venus fluorescence into the FRET channel. Thirdly, cells expressing Cerulean alone were used to measure the spectral crosstalk parameters *R*_*D1*_, and *R*_*D2*_, corresponding to the bleed-through of Cerulean fluorescence into the FRET and Venus channels, respectively. Fourth, the instrument-specific calibration parameters *f*_*Ven*_*/f*_*Cer*_ and *g*_*Ven*_*/g*_*Cer*_, corresponding to the ratios of the fluorescence excitation and emission of Venus and Cerulean, respectively, were measured on the day of the experiment using Cerulean-Venus dimers with known FRET efficiencies. Fifthly, cells co-expressing Cerulean and Venus were used to estimate concentration-dependent collisional FRET. For each cell, spectral crosstalk was then accounted for with Eq 5 – 7.

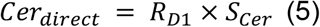

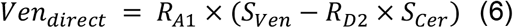

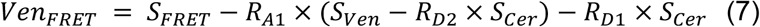

After unmixing the spectra, *f*_*A*_*/f*_*D*_ and *g*_*A*_*/g*_*D*_ were obtain by from the Cerulean-Venus dimer data by finding the slope and intercept of the linear relationship shown in Eq 8.

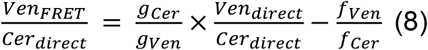

Donor-centric FRET efficiencies (E_D_) were then calculated with Eq 9.

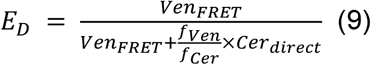

The relative proportion of Cerulean (N_Cer_) and Venus (N_Ven_) fluorophores in each cell were then determined from Eq 10 and 11, respectively.

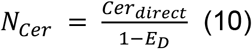

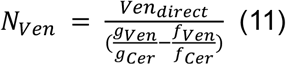

A 1:1 binding isotherm was then imposed by iteratively fitting the data to Eq 12,

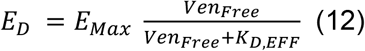

where *Ven*_*Free*_ is determined with Eq 13,

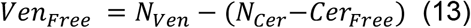

in which *Cer*_*Free*_ is computed from Eq 14.

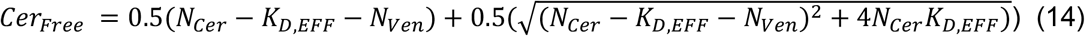

For each FRET pair, *K*_*D,EFF*_, *E*_*Max*_, and 95% confidence intervals were obtained through iterative constrained least squares analysis. Data were obtained from two independent transfections.

### Phylogenetic tree construction

The sequences of the nine human Na_V_ CTDs were obtained from UniProt^95^. The following residues were used for each isoform: Na_V_1.1: 1787-1954; Na_V_1.2: 1777-1944; Na_V_1.3: 1772-1939; Na_V_1.4: 1599-1776; Na_V_1.5: 1773-1940; Na_V_1.6: 1767-1934; Na_V_1.7: 1761-1928; Na_V_1.8: 1723-1890; Na_V_1.9: 1605-1772. The sequences were aligned with the BlastP algorithm and the phylogenetic tree constructed using the BLAST server^96^.

### Adeno-associated virus generation

Adeno-associated virus for the expression of ELIXIR, and control virus for the expression of GFP, were purchased from VectorBuilder. To monitor the expression of ELIXIR it was cloned with a C-terminal P2A site followed by mCherry (ELIXIR-P2A-mCherry).

### Quantification of ELIXIR-cpp uptake by aMVMs

Uptake of the ELIXIR-cpp was monitored using the same peptide as used in the fluorescence anisotropy experiments. Freshly dissociated aMVM were plated in a 12 well plate and incubated with different concentrations (100 nM – 10 μM) of ELIXIR-cpp for 2 hours at 37 °C in Tyrode’s solution (135 mM NaCl, 4 mM KCl, 1 mM MgCl_2_, 10 mM HEPES, 1 mM CaCl_2_ (pH 7.4 using NaOH)). Cellular uptake was monitored with an LSR II (BD Biosciences) flow cytometer equipped with 405 nm, 488 nm, and 633 nm lasers for excitation and 18 different emission channels as previously described^26^. Forward and side scatter signals were detected and used to gate for single cells that were living. Flow cytometric signals were collected at a medium flow rate (2k - 8k events/sec). Histograms corresponding to the single-cell population of the fluorescein channel were analyzed using custom MATLAB software (Mathworks).

### Brain slice electrophysiology

Congenic C57BL/6J mice heterozygous for the *SCN8A*-N1768D allele were crossed with each other to generate experimental *SCN8A*^N1768D/N1768D^ mice and wild-type *SCN8A*^+/+^ littermate controls. Mice were kept in the vivarium on a 12-h light-dark cycle. Mice had access to regular chow and water, ad libitum. Both male and female mice were used. Mice were bred and procedures were conducted at the Columbia University Institute of Comparative Medicine, which is fully accredited by the Association for Assessment and Accreditation of Laboratory Animal Care and were approved by the Columbia Institutional Animal Care and Use Committee (protocol #AC-AABI1551).

Intracranial injection of adenovirus containing ELIXIR-P2A-mCherry was performed at P0 or P1. Pups were injected under hypothermic anesthesia. Bilateral intracerebroventricular injections of virus (1 µL/hemisphere) were performed using a Hamilton syringe to induce widespread expression in the cortex. Pups were quickly rewarmed and returned to their home cage following injection.

Slice electrophysiology was conducted at P18-P22, as homozygous expression of the *SCN8A*-N1768D allele causes early mortality. Animals were deeply anesthetized with isofluorane and decapitated, and the brain was quickly dissected into ice-cold high-sucrose slicing solution (26 mM NaHCO3, 2.5 mM KCl, 1.25 mM NaH2PO4, 10 mM MgCl2, and 0.5 mM CaCl2, 11 mM d-glucose, 234 mM sucrose, pH 7.3-7.4). 250 µm slices were cut using a Leica VT1500 Vibratome. Slices were transferred to 37°C aCSF (126 mM NaCl, 26 mM NaHCO3, 2.5 mM KCl, 1.25 mM NaH2PO4, 1.0 mM MgCl2, and 2.0 mM CaCl2, 10 mM d-glucose, pH 7.3-7.4) and incubated for 1 hour. All solutions were continuously bubbled with 95% O2 and 5% CO2. Individual slices were transferred to a recording chamber located on an upright microscope (BX51WI; Olympus) and were perfused with heated (32 °C) oxygenated aCSF.

Whole-cell current-clamp recordings were obtained using Multiclamp 700B and Clampex 10.7 software (Molecular Devices). Layer V pyramidal cells were targeted for patch-clamp experiments based on location, size, and shape. Cells expressing ELIXIR-P2A-mCherry were identified using a fluorescent light source (Olympus). Both ELIXIR-P2A-mCherry expressing and non-expressing cells were targeted for patch clamp to generate within-subject control data. Intracellular solution contained 127 mM K-gluconate, 10 mM HEPES, 8 mM NaCl, 0.6 mM EGTA, 4 mM ATP, and 0.3 mM GTP, pH 7.2. When patch electrodes were filled with intracellular solution, their resistance ranged from 3.5 to 6 MΩ. Access resistance was monitored continuously for each cell and any cell that changed by more than 15% was excluded from analyses.

Holding current was delivered to adjust resting membrane potential to -75 mV. AP-generating properties of neurons were tested by injecting 1 sec square pulses of depolarizing current incrementing in 20 pA steps, up to 1 nA. AP threshold was defined as the membrane voltage at which the slope crossed 5 mV/ms. APs were automatically identified using this slope threshold and visually confirmed. APD_90_ was defined as the time during which the membrane voltage exceeded 10% of the peak amplitude. Half-width was defined as the time during which the membrane voltage exceeded 50% of the peak amplitude. Maximum rise rate was defined as the maximum of the first derivative of the rising phase of the AP. Maximum fall rate was defined as the minimum of the first derivative of the falling phase of the AP. Depolarization block threshold was defined as the current step at which the number of APs produced fell to less than ½ of the maximum. Brain slice electrophysiology data was analyzed offline with custom MATLAB software.

### Enzymatic isolation of ventricular CaMKIIδ_C,OE_ cardiomyocytes

Young adult (8 to 10-wk-old, male and female) CaMKIIdδ_C,OE_ mice and WT littermate control mice in the C57BL6/J background were used. Mice were kept at standard temperature, humidity, and lighting. Food (Teklad, 2018) and fresh, sterile drinking water were provided *ad libitum*. Before terminal surgery, mice were injected with heparin (400 U/kg body weight) and anesthetized with isoflurane (5% in an induction chamber, then 1.5 - 3% via nose cone). Hearts were excised and retrograde perfused on a constant flow Langendorff apparatus (4 min, 37 °C) with Ca^2+^-free normal Tyrode’s solution, gassed with 100% O_2_. The heart was then perfused for 12 - 17 min with 100 mg collagenase (type 2, Worthington Biochemical Corp) and 1.4 mg protease (type XIV, Sigma-Aldrich) in 50 mL of Tyrode’s solution (with 10 μmol/L Ca^2+^) to enzymatically isolate cardiomyocytes. Following digestion, the myocytes were gently triturated with a pipette, then filtered through a nylon mesh and allowed to sediment for ∼ 10 min. The sedimentation was repeated three times with increasing [Ca^2+^] from 0.125 to 0.25 to 0.5 mmol/L. Finally, ventricular myocytes were kept in Tyrode’s solution (0.5 mmol/L Ca^2+^) at room temperature until use. All animal handling procedures were in strict compliance with an approved protocol (#23175) of the Institutional Animal Care and Use Committee at the University of California, Davis conforming to the NIH Guide for the Care and Use of Laboratory Animals (8th edition, 2011).

### CaMKIIδ_C,OE_ cardiomyocyte electrophysiology

Isolated single murine ventricular cardiomyocytes were placed in a temperature-controlled perfusion chamber (Warner Instruments) and mounted on a Leica DMI3000 B inverted microscope (Leica Microsystems). Cells were bathed at 37°C, for 10 minutes, before starting, and continuously perfused (2 mL/min) during experiments with Tyrode’s solution containing (in mmol/L): NaCl 140, KCl 4, CaCl_2_ 1.8, MgCl_2_ 1, HEPES 5, Na-HEPES 5, glucose 5.5; pH=7.4. Electrodes were fabricated from borosilicate glass (World Precision Instruments) and had tip resistances of 2 to 3 MΩ when filled with internal solution which contained (in mmol/L): K-aspartate 100, KCl 30, NaCl 8, Mg-ATP 5, phosphocreatine dipotassium salt 10, HEPES 10, EGTA 0.01, cAMP 0.002, and calmodulin 0.0001; pH=7.2 (with KOH). Using this internal solution, the intracellular Ca^2+^ transient and contraction of the cardiomyocyte are preserved. An Axopatch 200B amplifier (Axon Instruments Inc.) was used for recording and signals were digitized at 50 kHz by a Digidata 1322A A/D converter (Axon Instruments) under software control (pClamp10.4). Series resistance was typically 3 to 5 MΩ and was compensated by ≥90%. Experiments were discarded when the series resistance was high or increased by ≥20% during the recordings. All experiments were conducted at 37±0.1°C.

APs were recorded in whole-cell current-clamp conditions where cells were stimulated with supra-threshold depolarizing pulses (2 ms duration) delivered via the patch pipette at 1 Hz. AP duration at 90% repolarization (APD_90_) was used to characterize AP repolarization. Series of 50 consecutive APs were analyzed to estimate short-term variability of the APD_90_. Short-term variability was calculated with equation 5,

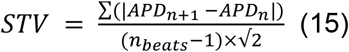

where APD_n_ and APD_n+1_ indicate the durations of the n^th^ and (n+1)^th^ AP, and n_beats_ denotes the total number of consecutive beats analyzed.

For voltage-clamp late Na^+^ current measurements, internal solution contained (in mmol/L): CsCl 110, tetraethylammonium chloride 20, Mg-ATP 5, HEPES 10, phosphocreatine disodium salt 5, calmodulin 0.0001, EGTA 10, CaCl_2_ 4.1 (free [Ca^2+^] =100 nmol/L), pH=7.20. Bath solution contained (in mmol/L): NaCl 140, CsCl 4, CaCl_2_ 1.8, MgCl_2_ 1, HEPES 5, Na-HEPES 5, glucose 5.5, 4-aminopyridine 5, nifedipine 0.01, pH=7.4. The late Na^+^ current magnitude was measured at the end of a 500 ms depolarizing pulse from a holding potential of -120 mV to -40 mV. The current amplitude was normalized to the capacitance (late Na^+^ current density) of each cell. Chemicals and reagents were purchased from Sigma-Aldrich. A subset of cells was preincubated with ELIXIR-cpp (3 μmol/L, 4 hours).

### Adeno-associated virus infection of Na_V_1.5 F1759A mice and isolation of cardiomyocytes

All procedures conducted with the Na_V_1.5 F1759A mice strictly adhered to a protocol approved by Institutional Animal Care and Use Committee at Columbia University. The Na_V_1.5 F1759A mouse line was previously generated by breeding Na_V_1.5 F1759A mice, in a B6CBA/F2 hybrid background, with cardiac rtTA mice in a FVB/N background to generate doxycycline-inducible transgenic mice^2^. Mice were kept on a 12-hour light-dark cycle, and fresh food and water were provided *ad libitum*. Both male and female mice were used. At postnatal day 4, Na_V_1.5 F1759A mouse pups were administered 1.03×10^11^ GC/g of adeno-associated virus 9 encoding either GFP or ELIXIR through a dorsal subcutaneous injection. ECG and electrophysiology experiments were conducted eight weeks after the injection. Prior to experimentation, the expression of Na_V_1.5 F1759A was induced with doxycycline chow overnight. Ventricular myocytes were isolated as described above.

### Confocal imagining

Images of freshly dissociated myocytes were collected on a Nikon Ti Eclipse inverted microscope equipped with a Yokogawa CSU-X1 spinning disk confocal and a CMOS camera from Andor Zyla scientific using a 60x oil-immersion objective.

### ECG analysis

Six-lead electrocardiograms were recorded from isoflurane-anesthetized mice using an Emka ECG with subcutaneous electrodes and IOX software (Emka Technologies). The RR QRS and QT durations were measured manually with IOX software. The QT duration was corrected for heart rate with Mitchell’s formula^74^ (equation 16),

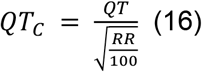

where QT and RR are the measured QT and RR interval, respectively.

### Differentiation of induced-pluripotent stem cell derived cardiomyocytes

Induced pluripotent stem cells were obtained and differentiated into cardiomyocytes as described previously^26^. Prior to differentiation human iPSCs were maintained in mTeSR medium (Stem Cell Technologies) and passaged every 4 - 6 days onto Matrigel-coated plates (Corning). On the first day of differentiation (day 0), embryoid bodies (EBs) were generated by treating the iPSCs with 1 mg/mL Collagenase B (Roche, Cat #: 11088807001) for 1 h, or until cells detached from the plate. The cells were then collected and centrifuged at 300 rcf for 3 min and resuspended as small clusters of 50–100 cells by gentle pipetting in differentiation medium (RPMI 1640 (Thermo Fisher Scientific, Cat #: 11875085) with 2 mM L-glutamine (Thermo Fisher Scientific, Cat #: 25030149), 0.4 mM monothioglycerol (Millipore Sigma, Cat #: M6145), and 50 μg/mL ascorbic acid (Millipore Sigma, Cat #: A4403)). Differentiation medium was supplemented with 2 ng/mL BMP4 (R&D Systems) and 10 μM Rock inhibitor (Y-27632 dihydrochloride, Tocris Fisher Cat #: 1254/50). EBs were cultured on ultra-low attachment 6-well plates (Corning Costar, Cat #: 3471) in a humidified incubator at 37°C, 5% CO_2_, and 5% O_2_. On day 1, the media was changed to differentiation media supplemented with 20 ng/mL BMP4 (R&D Systems), 20 ng/mL Activin A (R&D Systems), 5ng/mL bFGF (R&D Systems). On day 3, EBs were harvested and washed once with RPMI 1640. Medium was changed to differentiation media supplemented with 5 ng/mL VEGF (R&D Systems) and 5 μM XAV939 (Reprocell-Stemgent, Cat #: 04-0046). From this point on, every 2-3 days media was replaced with media supplemented with only 5 ng/mL VEGF (R&D Systems). Following differentiation, the cells were allowed to mature for 20 – 28 days before being transduced with ELIXIR or GFP AAV9. Experiments were conducted ∼5 days following transduction. Expression was visually confirmed by monitoring mCherry or GFP fluorescence.

### Statistical analysis

Pooled data are presented as the mean ± SEM. Statistical analysis was performed in Graphpad Prism 10.0.0. Normality was tested using a D’Agostino-Pearson normality test. For normally distributed data requiring multiple comparisons, a one-way analysis of variance (ANOVA) followed by Dunnett’s multiple comparisons was used. Multiple comparisons of nonnormally distributed data were done with a Kruskal-Wallis test followed by a Dunn’s post hoc test. Comparisons of two groups were conducted with a 2-tailed Student’s t-test (normally distributed data), or a Mann-Whitney U test (nonnormally distributed data). Differences with a P-value < 0.05 were considered statistically significant. For recordings conducted on aMVMs, the normality of the data was assessed by Shapiro-Wilk test and the equality of group variance was tested using the Brown-Forsythe test. Statistical significance of differences was determined using Mann-Whitney U test, and ANOVA with an appropriate multiple comparisons test, when applicable. Interaction between genotype and treatment was determined using a two-way ANOVA. Animals were grouped with no blinding but randomized in cellular experiments. Fully blinded analysis was not performed in cellular studies because the same person carried out the experiments and analysis. Group sizes were determined by an a priori power analysis with an α of 0.05 and power of 0.8, to detect a 20% signal difference at the endpoint.

### Images of protein structures

All protein structure images and vacuum electrostatics were generated with PyMOL (Schrödinger LLC). Unless otherwise stated, the Na_V_1.5_CTD_ is shown as gray cylinders and the ELIXIR peptide as a light blue (actinium) helix.

## Code availability

The peptide design protocol described here is available through Google Colab at: https://colab.research.google.com/github/sokrypton/ColabDesign/blob/v1.1.1/af/examples/peptide_binder_design.ipynb.

Matlab scripts used to analyze multichannel late current recordings are available on GitHub at: https://github.com/manubenjohny/LateCurrent.

## Acknowledgements

We would like to thank Audrey L. Kochiss for technical assistance, Dario Sirabella and Barbara Corneo from the Columbia University Stem Cell Core for assistance with iPSC-CMs differentiation, Jerry Chang from the Columbia University Precision Biomolecular Characterization Facility for fluorometer (NSF award 1828491) access, and the Columbia Center for Translation Immunology Flow Cytometry core which is supported in part by National Institutes of Health award S10RR027050. These studies were supported by funding from the National Institutes of Health to M.B.J. (R01 HL163576); R.M. (T32 HL120826); JK (K08 HL151969); S.O. (DP5OD026389); C.D.M. (R01 NS126392) and D.M.B. (P01-HL141084 and R01-HL142282) and from the American Heart Association to BH (23CDA1051603) and JK (CDA 35320208). We thank Dr. Michael Hammer (University of Arizona, Tucson, AZ) for the kind gift of the *SCN8A*-N1768D animal model. We thank Dr. Gordon Tomaselli for the kind of gift of the DKPQ hiPSC model. This content is solely the responsibility of the authors and does not necessarily represent the official views of the National Institutes of Health.

## Author contributions

R.M., and M.B-J. conceptualized and designed the project, S.O. developed the protocol for peptide hallucination, R.M., B.H., E.C., T.C., L.F., A.R., M.Y., N.C., and M.B-J. performed experiments; R.M., B.H., E.C., L.F., N.C., and M.B-J. analyzed data; R.M., E.C., B.H., R.J., E. W. S.O.M., J.K., C.D.M, D.M.B., and M.B-J. contributed reagents / analytic tools; S.O.M., S.O., J.K., C.D.M., D.M.B., and M.B-J. funding acquisition; R.M., B.H., and M.B-J made figures and wrote the original manuscript draft; and all authors revised the manuscript.

## Materials and Correspondence

All material requests and correspondence should be addressed to the corresponding author Manu Ben-Johny.

