## Supplementary Information for "De novo Design of a Peptide Modulator to Reverse Sodium Channel Dysfunction Linked to Cardiac Arrhythmias and Epilepsy"

**Supplemental Information**

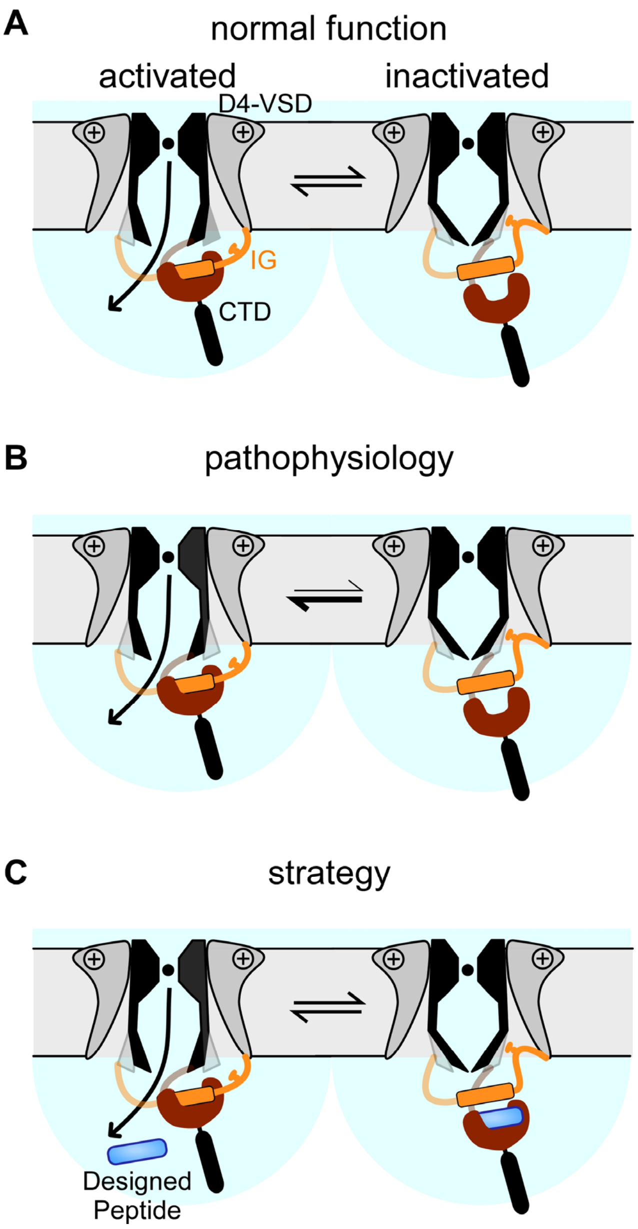


**Figure S1. Conceptual schematic of the mechanism of action of the designed peptide.**

**(A)** Following channel activation, the inactivation gate (IG) dissociates from the EF-hand like region (EFL, dark red) allowing it to interact with a pore-proximal site and inactivate $Na_V1.5$ (black/gray).

**(B)** In pathophysiology, IG has reduced efficacy for binding the pore-proximal site and remains associated to CTD, which allows for the continued flow of $Na^+$ into the cell.

**(C)** By competitively displacing the IG from the EFL, the designed peptide (ELIXIR) may promote channel inactivation and inhibit $I_{NaL}$.

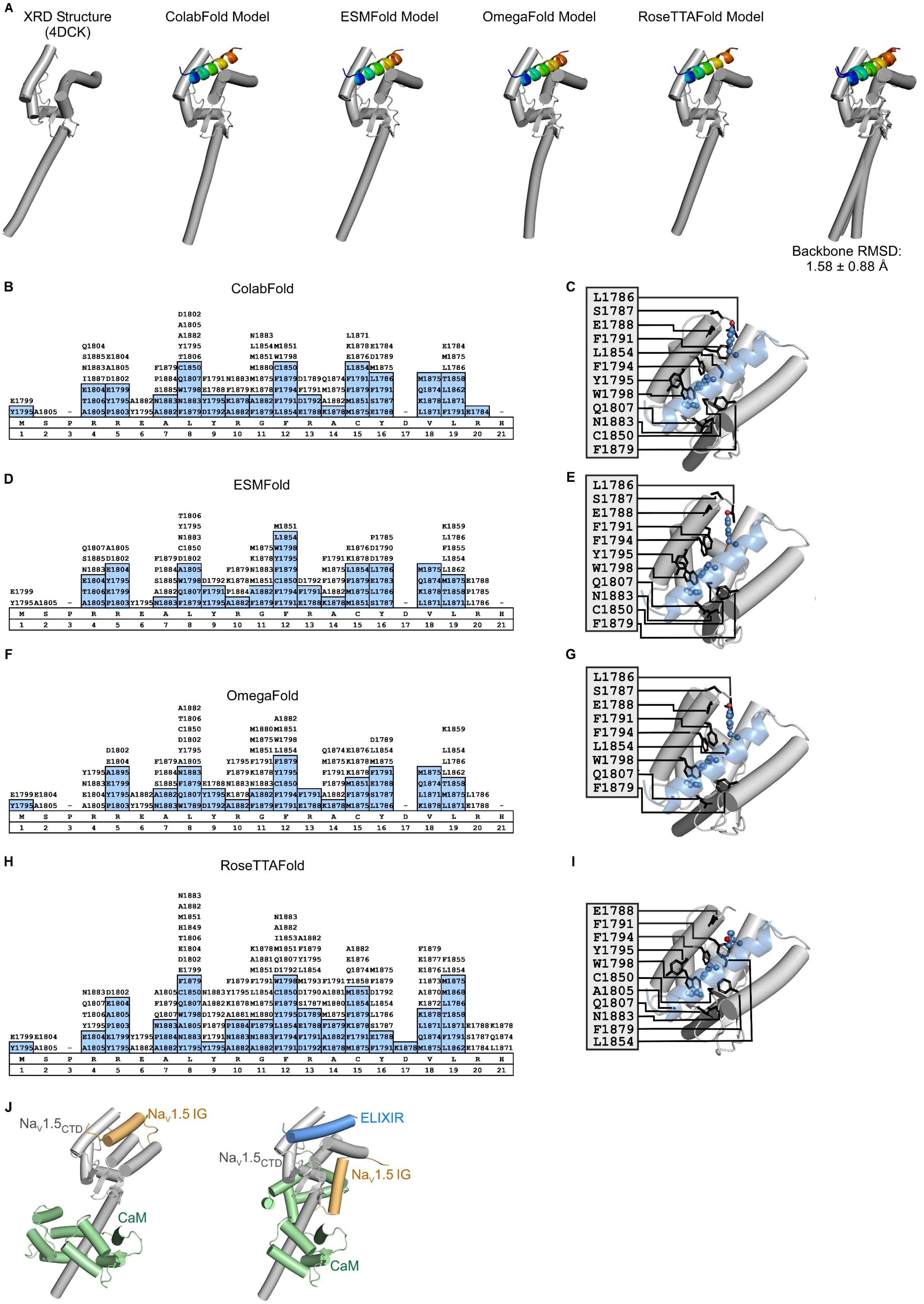
A
XRD Structure (4DCK)
ColabFold Model
ESMFold Model
OmegaFold Model
RoseTTAFold Model
Backbone RMSD: 1.58 ± 0.88 Å
B
ColabFold
C
L1786
S1787
E1788
F1791
L1854
F1794
Y1795
W1798
Q1807
N1883
C1850
F1879
D
ESMFold
E
L1786
S1787
E1788
F1791
F1794
Y1795
W1798
Q1807
N1883
C1850
F1879
F
OmegaFold
G
L1786
S1787
E1788
F1791
F1794
L1854
W1798
Q1807
F1879
H
RoseTTAFold
I
E1788
F1791
F1794
Y1795
W1798
C1850
A1805
Q1807
N1883
F1879
L1854
J
NaV1.5CTD
NaV1.5 IG
CaM
ELIXIR
NaV1.5CTD
NaV1.5 IG
CaM

**Figure S2. Models of the ELIXIR-$Na_V$1.5 CTD complex.**

**(A)** Comparison of a high-resolution structure of the $Na_V$1.5 CTD (PDB ID: 4DCK) to computational models of the $Na_V$1.5 CTD-ELIXIR complex generated with ColabFold, ESMFold, OmegaFold, and RoseTTAFold. The $Na_V$1.5 CTD is shown as gray cylinders and the ELIXIR peptide is shown in rainbow (blue, N-terminus; red, C-terminus). The models were aligned using residues 1786 – 1892 of the model generated with ColabFold.

**(B; D; F; H)** Contacts of Structural Units analysis of the ELIXIR-$Na_V$1.5 CTD models generated with ColabFold (**B**), ESMFold (**D**), OmegaFold (**E**) and RoseTTAFold (**H**). $Na_V$1.5 residues within 6 Å (within 4.5 Å colored blue) of the ELIXIR peptide are listed.

**(C; E; G; I)** Residues of the $Na_V$1.5 CTD (black sticks) within 6 Å of the ELIXIR peptide shown in the ELIXIR-$Na_V$1.5 CTD complexes generated with ColabFold (**C**), ESMFold (**E**) OmegaFold (**G**), RoseTTAFold (**I**). The $Na_V$1.5 CTD is shown as gray cylinders and ELIXIR as a light blue (actinium) $\alpha$-helix. Models were aligned as in **A**.

**(J)** ColabFold generated models of the $Na_V$1.5 CTD + $Na_V$1.5 IG + calmodulin (CaM) complex without (left) and with (right) ELIXIR. CaM is shown as pale green cylinders and the $Na_V$1.5 IG as a light orange cylinder, the $Na_V$1.5 CTD and ELIXIR are colored as in **C**. Models were aligned as in panel **A**.

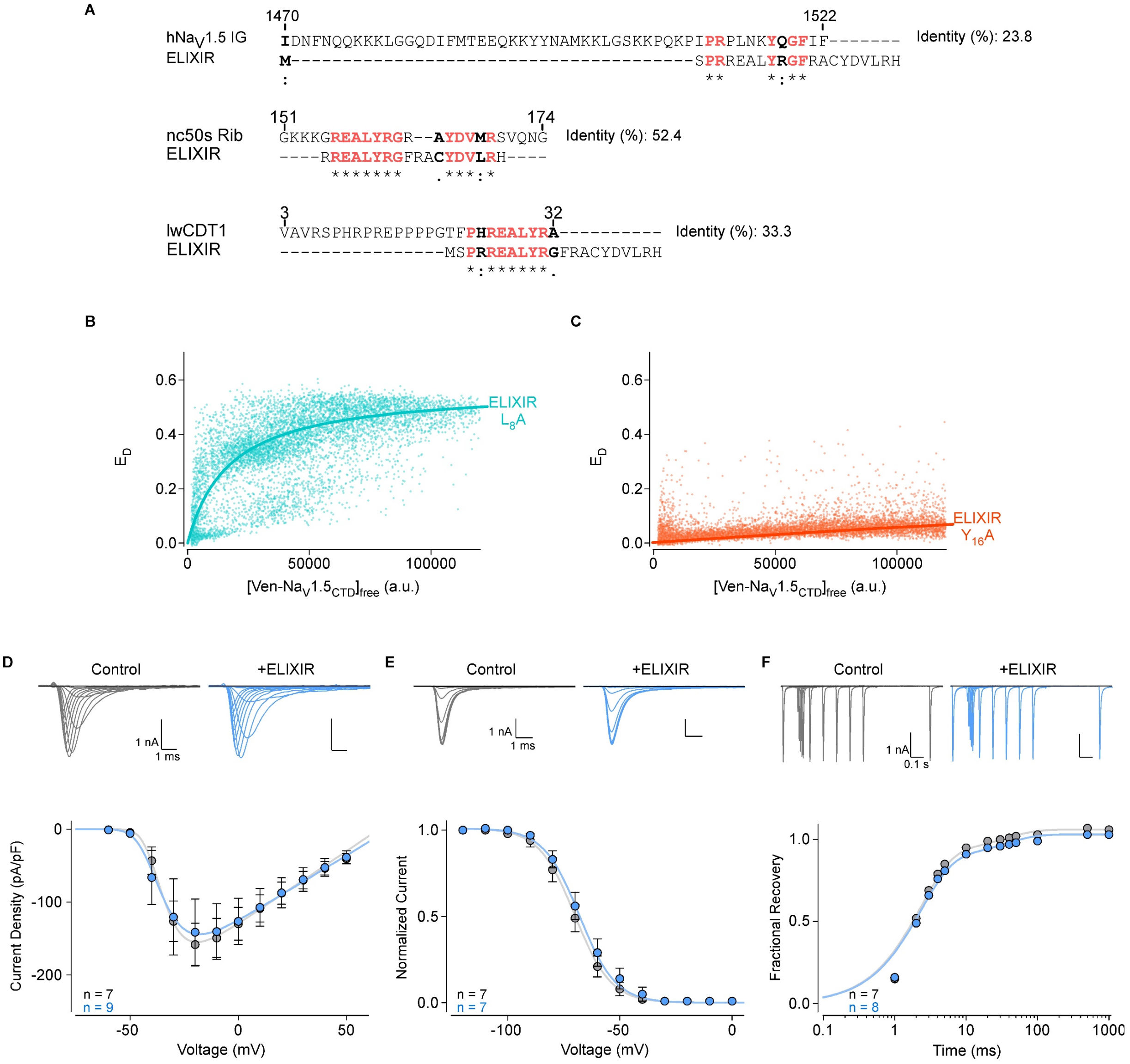


**G**

| | $V_{1/2}$ (mV) | $V_{rev}$ (mV) | Slope Factor | g |
|---|---|---|---|---|
| WT $Na_V1.5$ | -33.59 ± 1.52 | 69.48 ± 1.61 | 4.30 ± 0.53 | 33.83 ± 5.91 |
| WT $Na_V1.5$ + ELIXIR | -29.26 ± 2.76 | 69.51 ± 1.21 | 4.85 ± 0.39 | 36.20 ± 6.25 |

**Figure S3. ELIXIR sequence alignments.**

**(A )** Alignment of the human $Na_V1.5$ IG (top, $hNa_V1.5$ IG), *Nannochloropsis gaditana* 50s ribosome (middle**,** nc50s Rib) and *Leptonychotes weddellii* DNA replication factor CDT1 (bottom**,** lwCDT1) sequences with ELIXIR. Similar residues are shown in bold, and identical residues are colored red and shown in bold.

**(B-C)** FRET efficiency between Cerulean-tagged ELIXIR L8A (**B**) or ELIXIR Y16A (**C**) and Venus-tagged $Na_V1.5_{CTD}$ plotted against the concentration of free Venus-tagged $Na_V1.5_{CTD}$, a.u. is arbitrary units. The solid lines depict the fit of a 1:1 binding isotherm.

**(D)** Peak current density of wild-type $Na_V1.5$ in the absence (gray) or presence of ELIXIR (light blue). Data are shown as the mean ± SEM, with the number of replicate experiments (n) listed in parentheses. Exemplar currents are for wild-type $Na_V1.5$ alone (gray) and with ELIXIR (light blue) are shown above the plot.

**(E)** Steady state inactivation relationships for wild-type $Na_V1.5$ in the absence (gray) and presence of ELIXIR (light blue). Data are shown as the mean ± SEM, with the number of replicate experiments (n) shown in parentheses. Exemplar currents are for wild-type $Na_V1.5$ alone (gray) and with ELIXIR (light blue) are shown above the plot.

**(F)** Fractional recovery of wild-type $Na_V1.5$ from inactivation in the absence (gray) and presence of ELIXIR (light blue). Data are shown as the mean ± SEM, with the number of replicate experiments (n) listed. Exemplar currents are for wild-type $Na_V1.5$ alone (gray) and with ELIXIR (light blue) are shown above the plot.

**(G)** Summary of fit Parameters from whole-cell recordings shows minimal changes in voltage-dependence of channel activation and inactivation.

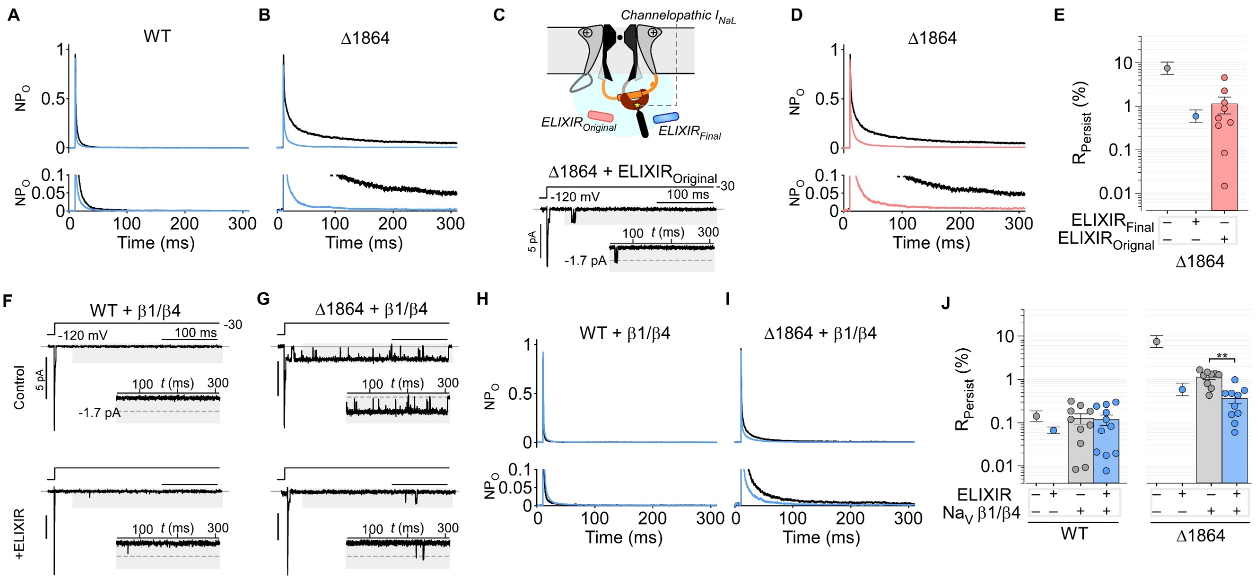


### Figure S4. Effect of Na$_V$ β-subunits and mutations on ELIXIR

**(A – B)** Comparison of the ensemble average NP$_o$ of Na$_V$1.5 wild-type (**A**) and Δ1864 (**B**) in the absence (black) or presence (light blue) of ELIXIR. The lower panel shows an expanded view near the baseline.

**(C)** Schematic of Na$_V$1.5 in the open state with ELIXIR$_{Original}$ (light red) and ELIXIR$_{final}$ (light blue) (top). Exemplar multichannel recording of Na$_V$1.5 Δ1864 co-expressed with with the initial ELIXIR peptide sequence (ELIXIR$_{Original}$; MSPRREALYRGFRACYDVLRH) (bottom). Inset shows the late phase enlarged for better visualization.

**(D)** Comparison of the ensemble average NP$_o$ of Na$_V$1.5 Δ1864 in the absence (black) and presence of ELIXIR$_{Original}$ (light red). The lower panel shows an expanded view near the baseline.

**(E)** Bar graph summarizing the effect of ELIXIR$_{Original}$ on the $I_{NaL}$ of Na$_V$1.5 Δ1864 (n = 9, 632 sweeps). Data are represented as mean ± SEM. The gray and light blue circles show the average (± SEM) R$_{persist}$ of the channel alone or with ELIXIR, respectively.

**(F – G)** Exemplar multichannel recordings of Na$_V$1.5 wild-type (**C**) and Δ1864 (**D**) in the absence (top) or presence (bottom) of ELIXIR. Insets show the late phase enlarged for better visualization.

**(H – I)** Comparison of the ensemble average NP$_o$ of Na$_V$1.5 wild-type (**E**) or Δ1864 (**F**) co-expressed with β1 and β4 in the absence (black) or presence (light blue) of ELIXIR. The lower panel shows an expanded view near the baseline.

**(J)** Bar graph summarizes effect of β1/β4-subunits and ELIXIR on $I_{NaL}$ of Na$_V$1.5 wild-type and Δ1864. Each bar, mean ± SEM. Na$_V$1.5 with β1/β4 alone (n = 10, 953 sweeps) or with β1/β4 and ELIXIR (n = 11; 921 sweeps), Na$_V$ Δ1864 with β1/β4 alone (n = 9, 775 sweeps) or with β1/β4 and ELIXIR (n = 10; 960 sweeps). **$p$ = 0.0021 (Na$_V$1.5 Δ1864+β1/β4+ELIXIR) compared to Na$_V$1.5 Δ1864+β1/β4 by a Brown-Forsythe test followed by Dunn's multiple comparison test.

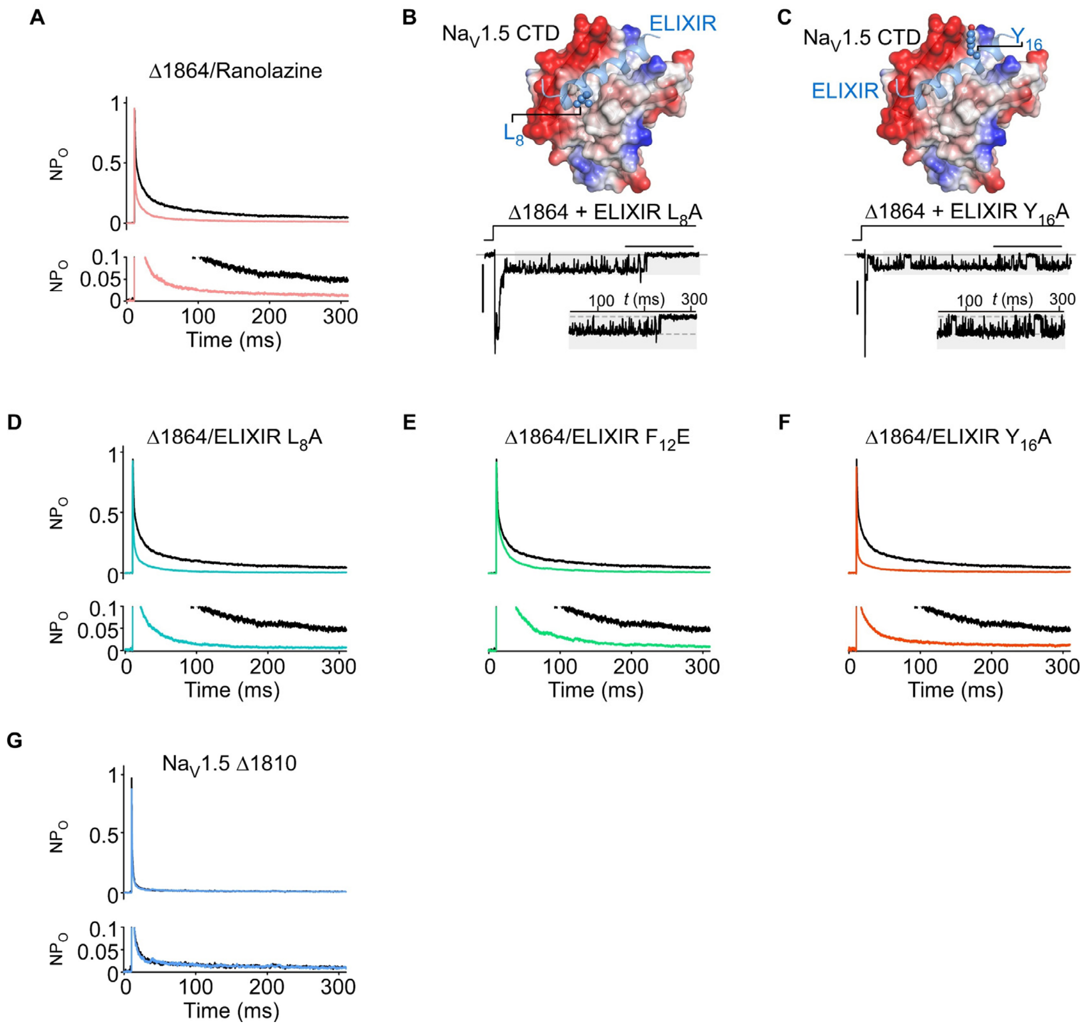


**Figure S5. Mutations in ELIXIR reduce $I_{NaL}$ inhibition.**

**(A)** Comparison of the ensemble average $NP_o$ of $Na_V1.5$ Δ1864 in the absence (black) and presence (pink) of 10 μM ranolazine. The lower panel shows an expanded view near the baseline.

**(B – C)** Top: vacuum electrostatic surface (positive; blue, neutral, white; negative, red) of the $Na_V1.5$ CTD EFL bound to ELIXIR with L8 (**I**) or Y16 (**J**) shown as ball-and-stick (top). Bottom: exemplar multichannel recordings of $Na_V1.5$ Δ1864 in presence of ELIXIR L8A (**I**) or ELIXIR Y16A (**J**). Insets show the late phase enlarged for better visualization.

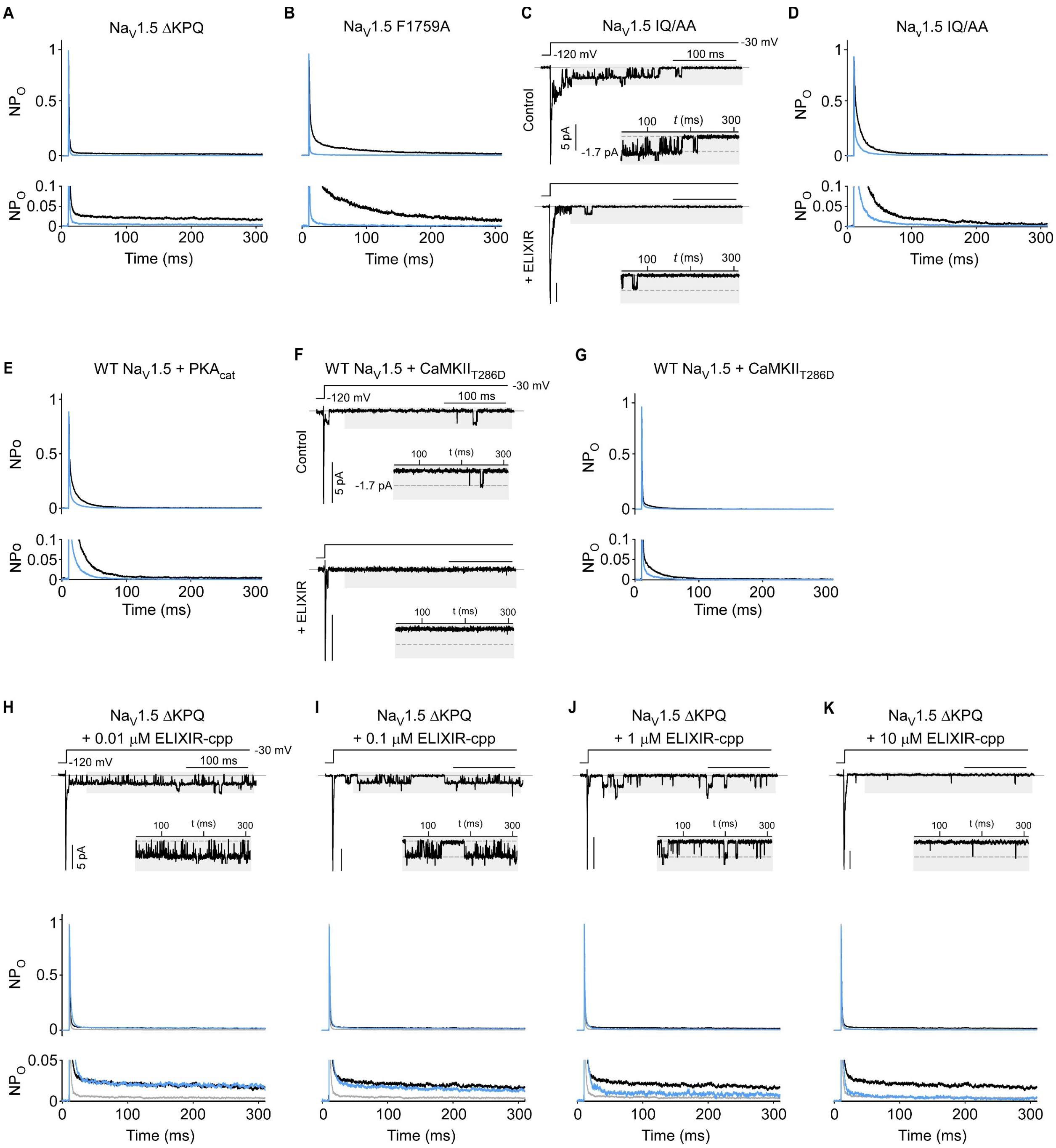

A
NaV1.5 ΔKPQ
NPO
Time (ms)
B
NaV1.5 F1759A
C
NaV1.5 IQ/AA
-120 mV
-30 mV
100 ms
Control
5 pA
-1.7 pA
t (ms)
+ ELIXIR
D
NaV1.5 IQ/AA
E
WT NaV1.5 + PKAcat
NPo
F
WT NaV1.5 + CaMKIIT286D
G
WT NaV1.5 + CaMKIIT286D
H
NaV1.5 ΔKPQ
+ 0.01 μM ELIXIR-cpp
I
NaV1.5 ΔKPQ
+ 0.1 μM ELIXIR-cpp
J
NaV1.5 ΔKPQ
+ 1 μM ELIXIR-cpp
K
NaV1.5 ΔKPQ
+ 10 μM ELIXIR-cpp

**Figure S6. Generality of $I_{NaL}$ inhibition by ELIXIR**

**(A - B)** Comparison of the ensemble average $NP_o$ of $Na_V$1.5 ΔKPQ (**A**) and $Na_V$1.5 F1759A (**B**) in the absence (black) and presence of ELIXIR (light blue). The lower panel shows an expanded view near the baseline.

**(C)** Exemplar multichannel recordings of $Na_V$1.5 IQ/AA in the absence (top) and presence (bottom) of ELIXIR. Insets show the late phase enlarged for better visualization.

**(D - E)** Comparison of the ensemble average $NP_o$ of $Na_V$1.5 IQ/AA (**D**) and WT $Na_V$1.5+$PKA_{Cat}$ (**E**) in the absence (black) and presence of ELIXIR (light blue). The lower panel shows an expanded view near the baseline.

**(F)** Exemplar multichannel recordings of WT $Na_V$1.5+$CaMKII_{T286D}$ in the absence (top) and presence (bottom) of ELIXIR. Insets show the late phase enlarged for better visualization.

**(G)** Comparison of the ensemble average $NP_o$ of WT $Na_V$1.5+$CaMKII_{T286D}$ (**G**) in the absence (black) and presence of ELIXIR (light blue). The lower panel shows an expanded view near the baseline.

**(H - K)** Exemplar multichannel recordings (top) and comparison of the ensemble average $NP_O$ (bottom) of $Na_V$1.5 ΔKPQ following treatment with 0.01 μM ELIXIR-cpp (**H**), 0.1 μM ELIXIR-cpp (**I**), 1 μM ELIXIR-cpp (**J**) and 10 μM ELIXIR-cpp (**K**). In each panel the ensemble average $NP_o$ of $Na_V$1.5 ΔKPQ in the absence of ELIXIR is shown in black, with ELIXIR overexpressed in gray and following treating with the designated concentration of ELIXIR-cpp in light blue. In the upper panel the insets show the late phase enlarged for better visualization. The bottom panel shows an expanded view, near the baseline, of the ensemble average $NP_O$.

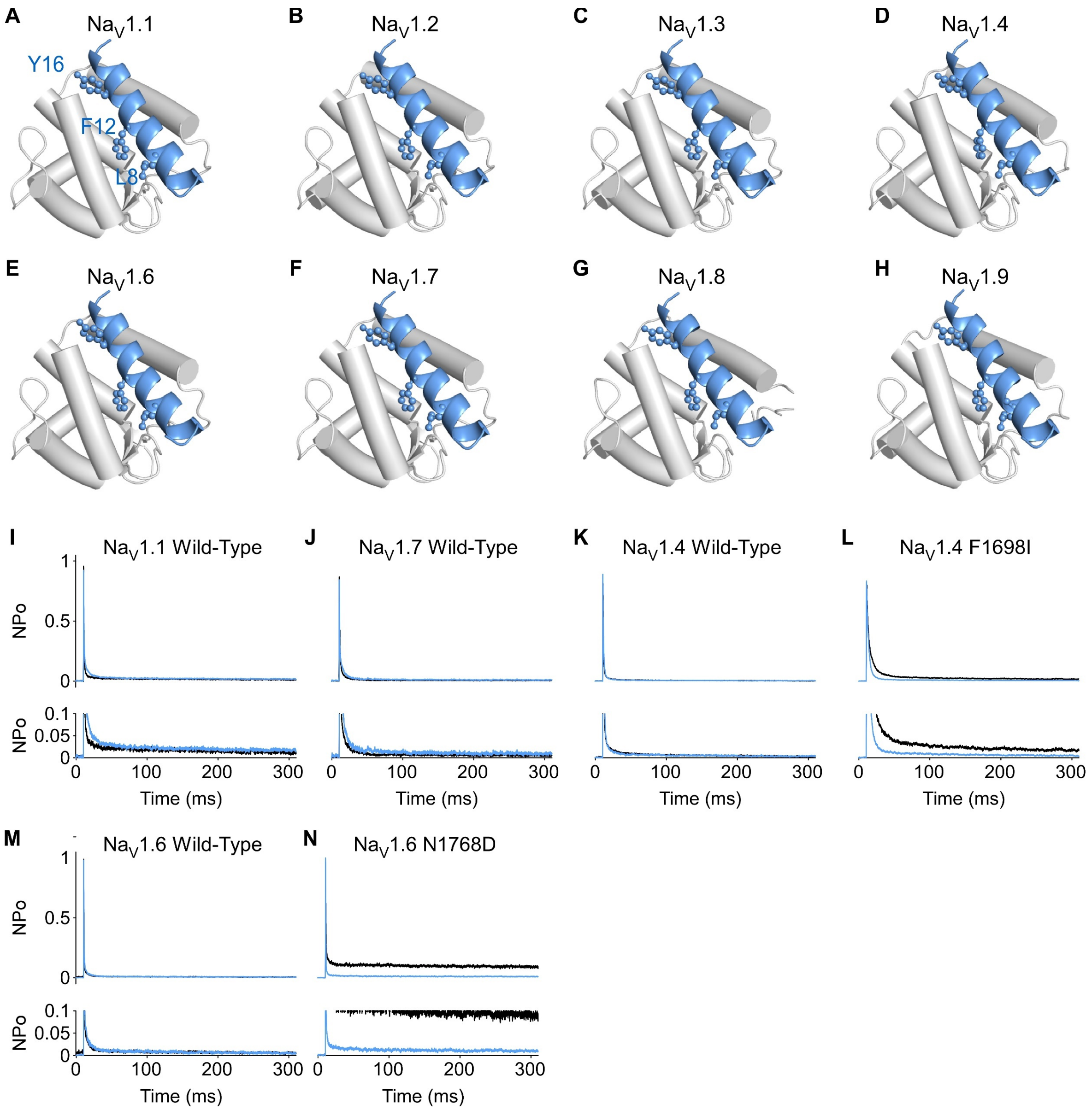


**Figure S7. ColabFold models of ELIXIR-Na$_V$ CTD complexes.**

**(A – H)** ColabFold generated models of ELIXIR (light blue) bound to the CTD (gray cylinders) of Na$_V$1.1 (**A**), Na$_V$1.2 (**B**), Na$_V$1.3 (**C**), Na$_V$1.4 (**D**), Na$_V$1.6 (**E**), Na$_V$1.7 (**F**), Na$_V$1.8 (**G**), and Na$_V$1.9 (**H**). ELIXIR residues L8, F12, and Y16 are shown as ball and stick. The structures were aligned to residues 1788 – 1893 of Na$_V$1.5. For clarity only the EFL region (residues 1788-1893 of Na$_V$1.5) of each CTD is shown.

**(I – N)** Comparison of the ensemble average NP$_o$ of WT Na$_V$1.1 (I), WT Na$_V$1.7 (**J**), WT Na$_V$1.4 (**K**), Na$_V$1.4 F1698I (**L**), WT Na$_V$1.6 (**M**) and Na$_V$1.6 N1768D (**N**) in the absence (black) and presence of ELIXIR (light blue). The lower panel shows an expanded view near the baseline.

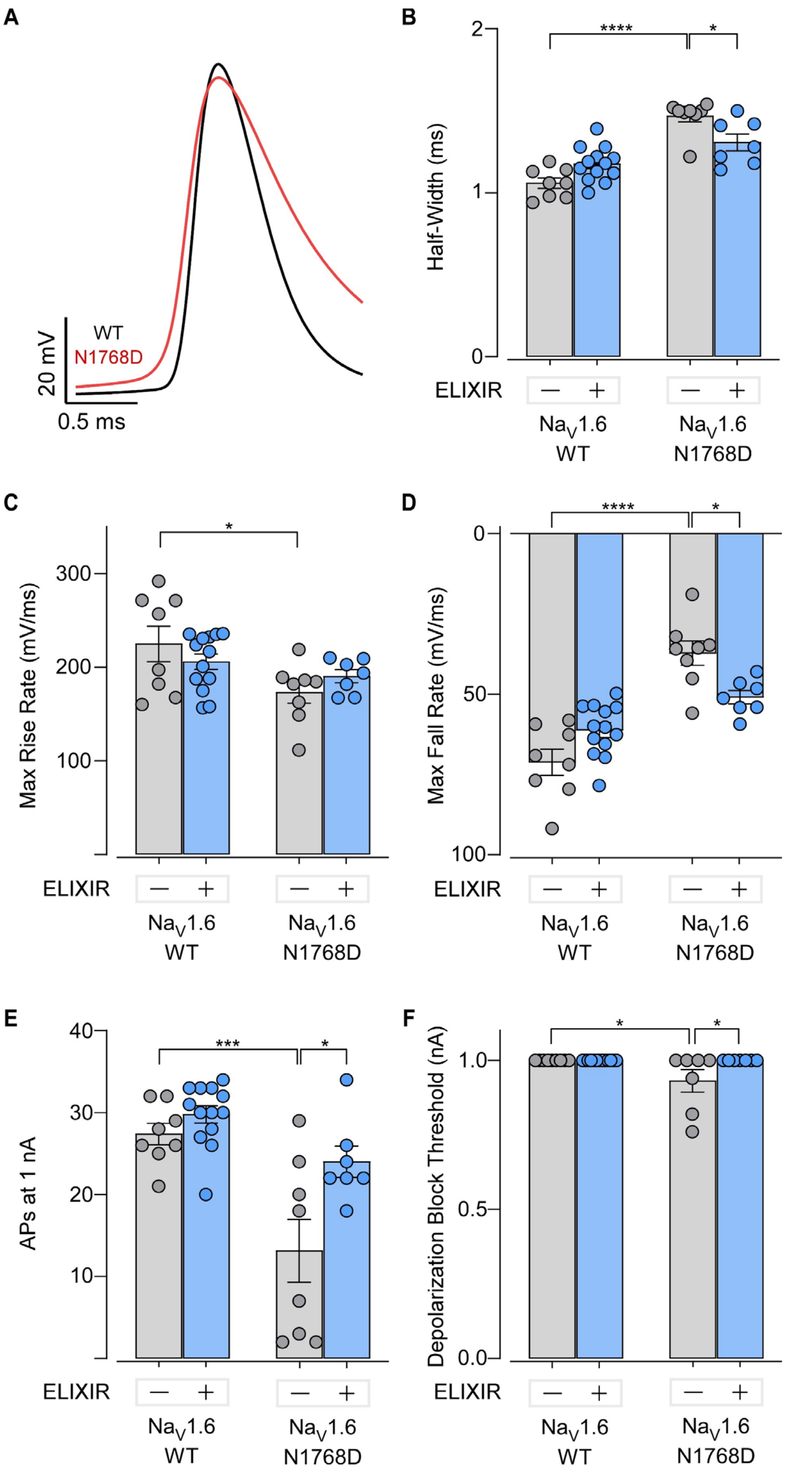

A
WT
N1768D
20 mV
0.5 ms
B
Half-Width (ms)
****
*
ELIXIR
Na$_V$1.6 WT
Na$_V$1.6 N1768D
C
Max Rise Rate (mV/ms)
*
D
Max Fall Rate (mV/ms)
****
*
E
APs at 1 nA
***
*
F
Depolarization Block Threshold (nA)
*
*

**Figure S8. Effect of ELIXIR on WT $Na_V$1.6 and $Na_V$1.6 N1768D neuronal AP properties.**

**(A)** Exemplar AP recordings of control neurons from WT $Na_V$1.6 (black) and $Na_V$1.6 N1768D (red) mice.

**(B)** Bar graph summarizing the effect of ELIXIR on the AP half width of control and ELIXIR expressing neurons from WT $Na_V$1.6 (control n = 8, N = 3 mice; ELIXIR n = 13, N = 3 mice) or $Na_V$1.6 N1768D (control n = 8, N = 2 mice; ELIXIR n = 7, N = 2 mice) mice. Each bar, mean ± SEM. ****$p < 0.0001$ (WT $Na_V$1.6 control), *$p = 0.0330$ ($Na_V$1.6 N1768D ELIXIR) compared to $Na_V$1.6 N1768D control by a two-way ANOVA followed by a Tukey's multiple comparisons test.

**(C)** Bar graph summarizing the effect of ELIXIR on the maximum rise rate of APs from control and ELIXIR expressing neurons from WT $Na_V$1.6 (control n = 8, N = 3 mice; ELIXIR n = 13, N = 3 mice) and $Na_V$1.6 N1768D (control n = 8, N = 2 mice; ELIXIR n = 7, N = 2 mice) mice. Each bar, mean ± SEM. *$p = 0.0296$ (WT $Na_V$1.6 control) compared to $Na_V$1.6 N1768D control by a two-way ANOVA followed by a Tukey's multiple comparisons test.

**(D)** Bar graph summarizing the effect of ELIXIR on the maximum fall rate of APs from control and ELIXIR expressing neurons from WT $Na_V$1.6 (control n = 8, N = 3 mice; ELIXIR n = 13, N = 3 mice) and $Na_V$1.6 N1768D (control n = 8, N = 2 mice; ELIXIR n = 7, N = 2 mice) mice. Each bar, mean ± SEM. ****$p < 0.0001$ (WT $Na_V$1.6 control), *$p = 0.0320$ ($Na_V$1.6 N1768D ELIXIR) compared to $Na_V$1.6 N1768D control by a two-way ANOVA followed by a Tukey's multiple comparisons test.

**(E)** Bar graph summarizing the effect of ELIXIR on the number of APs fired at 1 nA from control and ELIXIR expressing neurons from WT $Na_V$1.6 (control n = 8, N = 3 mice; ELIXIR n = 13, N = 3 mice) and $Na_V$1.6 N1768D (control n = 8, N = 2 mice; ELIXIR n = 7, N = 2 mice) mice. Each bar, mean ± SEM. ***$p < 0.0004$ (WT $Na_V$1.6 control), *$p = 0.0104$ ($Na_V$1.6 N1768D ELIXIR) compared to $Na_V$1.6 N1768D control by a two-way ANOVA followed by a Tukey's multiple comparisons test.

**(F)** Bar graph summarizing the effect of ELIXIR on the depolarization block threshold of control and ELIXIR expressing neurons from WT $Na_V$1.6 (control n = 8, N = 3 mice; ELIXIR n = 13, N = 3 mice) and $Na_V$1.6 N1768D (control n = 8, N = 2 mice; ELIXIR n = 7, N = 2 mice) mice. Each bar, mean ± SEM. *$p < 0.01884$ (WT $Na_V$1.6 control), *$p = 0.0289$ ($Na_V$1.6 N1768D ELIXIR) compared to $Na_V$1.6 N1768D control by a two-way ANOVA followed by a Tukey's multiple comparisons test.

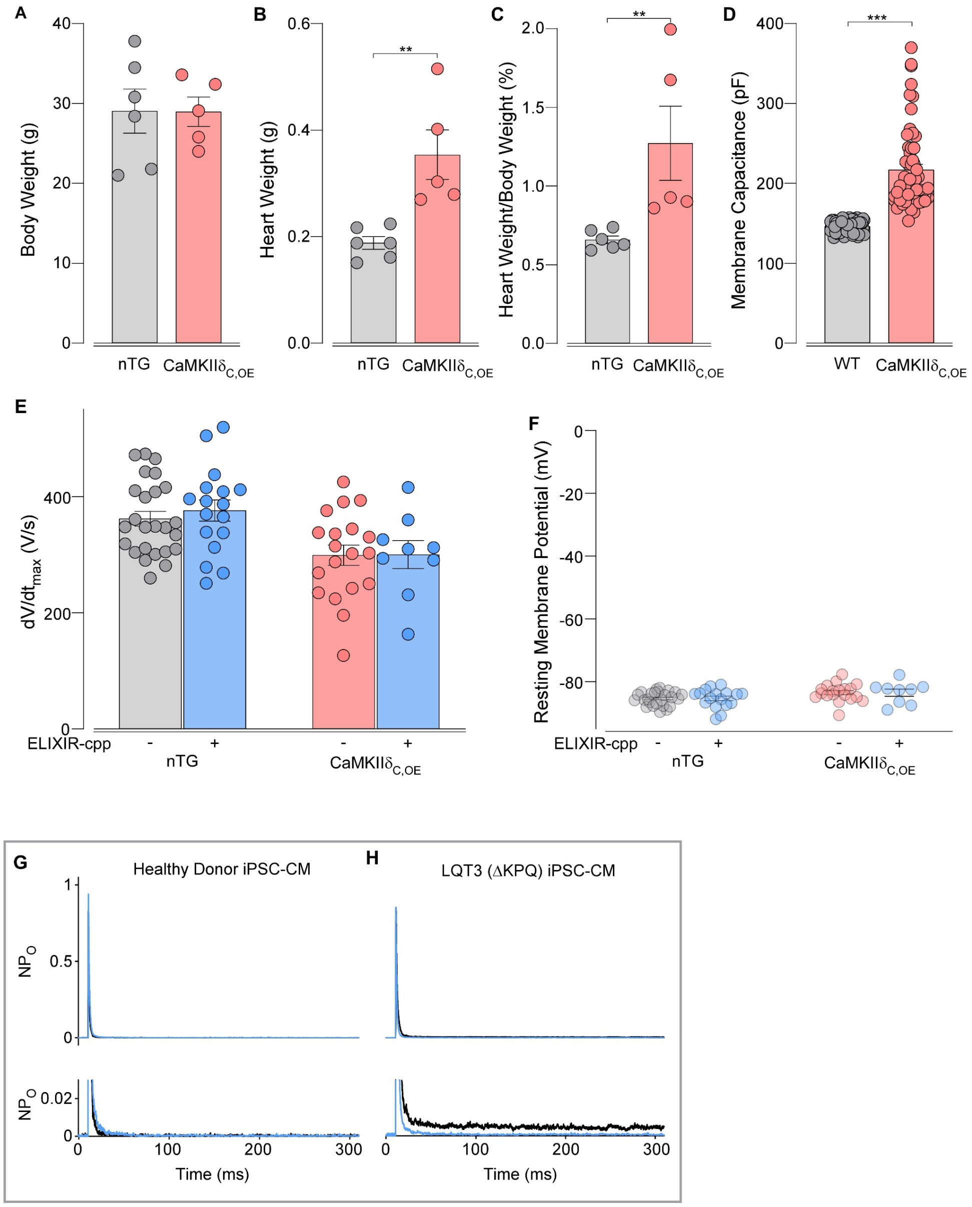


**Figure S9. Comparison of nTG and CaMKIIδ$_{C,OE}$ murine heart morphology**.

**(A – C)** bar graph comparison of the body weight (**A**), heart weight (**B**), and heart weight body weight ratio (**C**) of nTG (N = 6 mice) and CaMKIIδ$_{C,OE}$ (N = 5 mice) mice. Data are shown as mean ± SEM. ***p* = 0.004 for both heart weight and heart weight/body weight ratio by a Mann-Whiteny U test.

**(D)** bar graph comparison of the membrane capacitance of cardiomyocytes isolated from nTG (n = 80 cells, N = 6 mice) and CaMKIIδ$_{C,OE}$ (red, n = 56 cells, N = 5 mice) mice. *** *p* < 0.001 by a Mann-Whiteny U test.

**(E – F),** Comparison of the dv/dt (**E**) and resting membrane potential (**F**) of cardiomyocytes isolated from nTG mice in the absence (n = 25 cells, N = 6 mice) and presence (n = 17 cells, N = 6 mice) of ELIXIR-cpp or from CaMKII$\delta_{C,OE}$ mice in the absence (n = 19 cells, N = 5 mice) or presence (n = 9 cells, N = 4 mice) of +ELIXIR-cpp.

**(G – H)** Comparison of the ensemble average $NP_o$ of Healthy Donor (**G**), LQT3 (ΔKPQ) (**H**) iPSC-CMs, infected with adenovirus containing GFP (black) or ELIXIR (light blue). The lower panel shows an expanded view near the baseline.
